# mRNA- and factor-driven dynamic variability controls eIF4F-cap recognition for translation initiation

**DOI:** 10.1101/2021.06.17.448745

**Authors:** Burak Çetin, Seán E. O’Leary

## Abstract

mRNA 5ʹ cap recognition by eIF4F is a key step in eukaryotic translational control. While different mRNAs respond differently to eIF4F-directed regulation, the molecular basis for this variability remains unclear. We developed single-molecule fluorescence assays to directly observe eIF4F– mRNA interactions. We uncovered a complex interplay of mRNA features with factor activities that differentiates cap recognition between mRNAs. eIF4E–cap association rates are anticorrelated with mRNA length. eIF4A leverages ATP binding to differentially accelerate eIF4E– mRNA association; the extent of this acceleration correlates with translation efficiency *in vivo*. eIF4G lengthens eIF4E–cap binding to persist on the initiation timescale. The full eIF4F complex discriminates between mRNAs in an ATP-dependent manner. After eIF4F–mRNA binding, eIF4E is ejected from the cap by eIF4A ATP hydrolysis. Our results suggest features throughout mRNA coordinate in controlling cap recognition at the 5ʹ end, and suggest a model for how eIF4F–mRNA dynamics establish mRNA sensitivity to translational control processes.

## INTRODUCTION

Protein synthesis is a key stage of gene expression, and is highly regulated to maintain cellular viability (Hershey et al., 2012). In eukaryotes, translation is chiefly controlled during initiation. Translational control also allows gene expression to respond to changes in the cellular environment, and to external stimuli, on a timescale faster than transcriptional regulation (Hershey et al., 2012).

Translation initiation on most eukaryotic mRNAs involves the 5′ m^7^G(5ʹ)ppp(5ʹ)N cap (where N is the transcript +1 nucleotide) (Furuichi and Shatkin, 2000). Cap recognition by initiation factor eIF4F leads to recruitment of the small ribosomal subunit, in its 43S pre-initiation complex (PIC) (Fraser, 2015). The resulting 48S PIC is thought to scan linearly through the mRNA 5′ leader to the start codon (Kozak and Shatkin, 1978; Fraser, 2015; Hinnebusch and Lorsch, 2012). Cap recognition, PIC recruitment and start-codon selection are each major targets for translational control mechanisms (Jackson et al., 2010; Sonenberg and Hinnebusch, 2009).

mRNAs encoding different proteins are translated with widely differing efficiencies (Sonneveld et al., 2020; Genuth and Barna, 2018). Early kinetic modeling studies implicated variable cap recognition as crucial for “discrimination” between mRNAs competing for translation (Temple and Lodish, 1975; Godefroy-Colburn and Thach, 1981). Numerous subsequent studies have confirmed eIF4F as one key mediator of discrimination, and characterized the biology and biochemistry of cap recognition and its regulation (Gingras et al., 1999;Pelletier and Sonenberg, 2019; Kozak and Shatkin, 1978). mRNA features in the 5ʹ leader render translation more or less eIF4F-dependent (Hershey et al., 2012; Pelletier et al., 2015) – in particular, mRNAs with longer, more heavily-structured leaders show stronger dependence (Richter and Sonenberg, 2005). mRNA elements that define or modulate eIF4F dependence also contribute to the molecular mechanisms of pathogenesis in cancer, viral infection (Schneider and Mohr, 2003), developmental and neurodegenerative disorders (Amorim et al. 2018), and diet-induced obesity (Pelletier et al., 2015; Conn et al. 2021).

eIF4F is composed of three subunits. eIF4E binds the cap structure (Marcotrigiano et al., 1997; Siddiqui and Sonenberg, 2015; von der Haar et al., 2000; Hershey et al., 2012; Pelletier et al., 2015). eIF4G, a large, multidomain protein, binds eIF4E, mRNA, and eIF4A, a DEAD-box RNA helicase (Merrick, 2015; Altmann and Linder, 2010). eIF4G also directly contacts the 43S PIC (Altmann and Linder, 2010; Sonenberg and Hinnebusch, 2009). The cellular concentration of eIF4F is tightly regulated through eIF4E sequestration by 4E-binding proteins (4EBPs), which modulates translation in response to cell-signaling events (von der Haar et al., 2004; Richter and Sonenberg, 2005; Siddiqui and Sonenberg, 2015); yeast 4E-BPs are also phosphorylated, though the importance of this for translational control is less clear (Arndt *et al*., 2018; Zanchin and McCarthy, 1995).

Translation initiation is highly dynamic, proceeding through numerous intermediates with different molecular compositions and conformations on a seconds timescale (Prabhakar et al., 2019; Wang et al., 2019). Thus, understanding how eIF4F interacts with different mRNAs in real time during initiation, and how its subunits contribute to those interactions and their differences, is central to understanding mechanisms of mRNA discrimination and translational control. The advent of mRNA-based vaccines (Pardi et al., 2018; Kyriakidis et al, 2021) further underscores the need to understand eIF4F–mRNA interaction dynamics, since these interactions contribute significantly to translational output.

*In vivo* RIP-seq data from *Saccharomyces cerevisiae* indicate mRNA binding to eIF4E•eIF4G varies ∼7-fold, and, intriguingly, appears to anticorrelate with coding-sequence length (Costello et al., 2015; Thompson and Gilbert, 2017). This suggests mRNA features beyond the cap-proximal region, or even beyond the leader, may be important determinants of eIF4F–mRNA interaction. Indeed, an mRNA “length sensing” model has been proposed to contribute to variability in translation efficiency (Thompson and Gilbert, 2017). Features beyond the 5ʹ leader have also been proposed to contribute to the ability of eIF4A to promote PIC recruitment (Yourik et al., 2017). However, while the structural, thermodynamic, and kinetic bases of eIF4E interaction with the dinucleotide cap structure and capped oligoribonucleotides have been characterized (Marcotrigiano et al., 1997; Niedzwiecka et al., 2002; Slepenkov et al., 2008; Kaye et al., 2009; O’Leary et al., 2013; Carberry et al., 1989), information on real-time interactions with full-length mRNAs is limited.

We developed a suite of single-molecule fluorescence assays to directly observe real-time *Saccharomyces cerevisiae* eIF4F–mRNA recognition, and to dissect the contributions of the eIF4F subunits on individual, full-length yeast mRNAs. We applied these assays to define how the eIF4F complex dynamically coordinates with mRNA to control variability in cap-recognition efficiency between mRNAs.

## RESULTS

### mRNAs show a broad range of eIF4E binding interactions

From the ∼6,000 *S. cerevisiae* protein-coding genes (Hirschman et al., 2006) we chose mRNAs spanning a range of eIF4E binding in cells (Costello et al., 2015). *In vivo* mRNA enrichments in eIF4E and eIF4G are broadly similar; therefore, our selection also reflects enrichment in eIF4G. We restricted mRNA candidates to those that span a range of changes in translation efficiency (“TE”) on conditional knockdown of eIF4A, but where TE is not significantly dependent on the second translational helicase, Ded1p (Sen et al., 2015).

We arrived at four mRNAs that permute *in vivo* eIF4E binding and eIF4A translation dependence (Figure 1A) – with *in-vivo* eIF4E binding that is below average (*HXT2*), average (*NCE102*), or above average (*JJJ1, HSP30*), and where eIF4A knockdown either reduces translation efficiency (ΔTE < 0; *JJJ1, HXT2*), vs. where it remains unchanged or potentially increases (ΔTE > 0; *NCE102, HSP30*). To this set we added *MIM1* and *POP5*, to allow coverage over a broad range of coding-sequence lengths (0.3 – 1.7 kb). We transcribed, capped, and polyadenylated these mRNAs *in vitro* (Supplemental Table 1; Supplemental Figure 1A).

**Figure 1.**
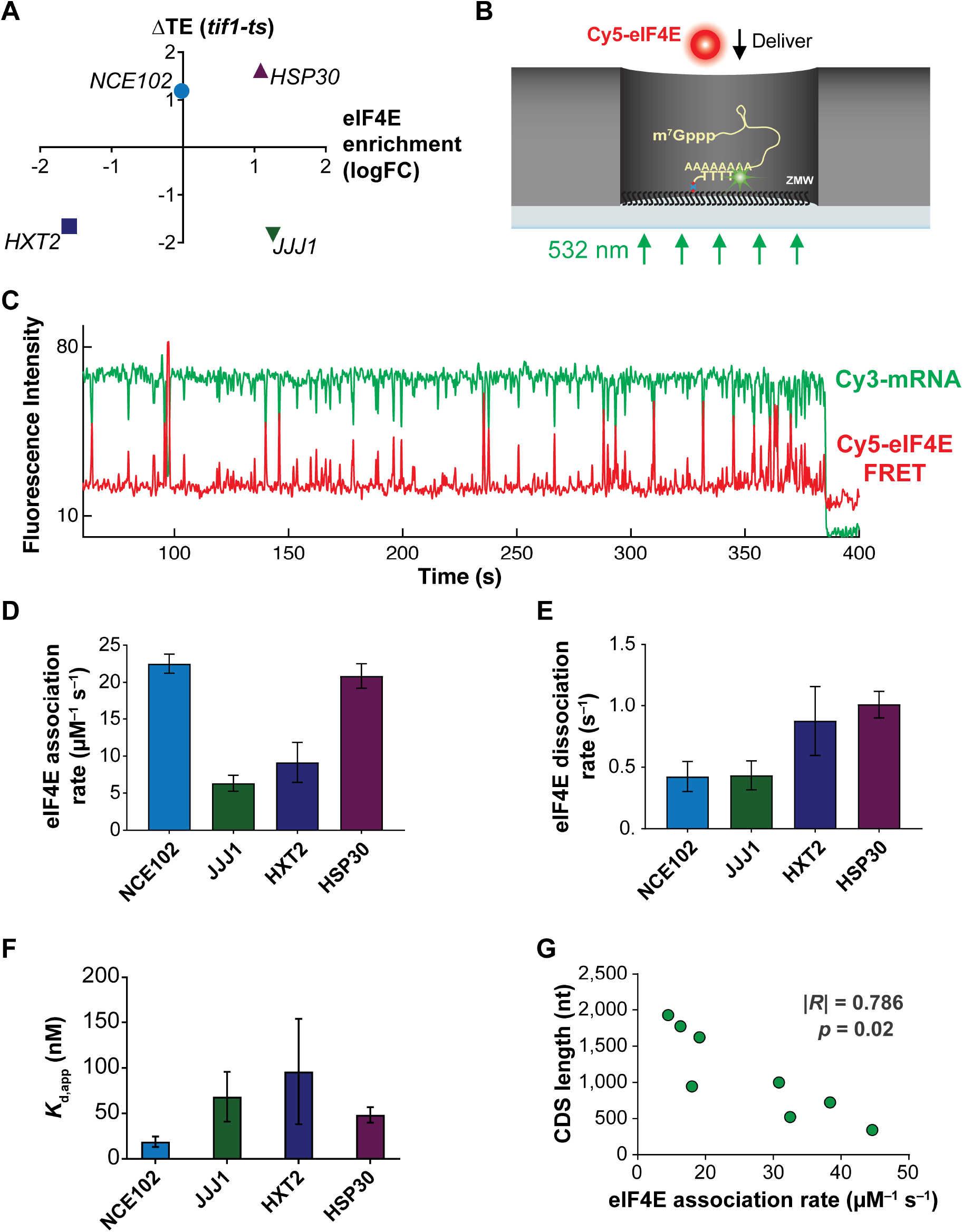
eIF4E interaction dynamics with full-length mRNAs depend on mRNA identity and length. **A.** Selection of mRNAs with varying *in-vivo* enrichment in eIF4E•eIF4G and translation dependence on eIF4A, as measured by Costello *et al*. (2015), and Sen *et al*. (2015). **B.** Schematic of single-molecule FRET experiment to detect binding of fluorescently-labeled eIF4E to surface-immobilized, fluorescently-labeled mRNA. **C.** Sample smFRET trajectory showing eIF4E–mRNA interaction in the absence of other eIF4F components. **D.** eIF4E–mRNA association rates. Error bars reflect the standard errors of the mean for three replicates of an experiment where the eIF4E–mRNA binding rate is measured across at least 100 mRNA molecules. **E.** eIF4E–mRNA dissociation rates from the experiments in D. **F.** eIF4E–mRNA equilibrium dissociation constants computed from the rates shown in D and E. **G.** Dependence of eIF4E–mRNA association rate on mRNA coding-sequence length. Note that the plot includes four previously published association rates from Çetin *et al*. (2020), to include sufficient data points for correlation.

We surface-immobilized the mRNAs for single-molecule fluorescence analysis of eIF4E binding, by hybridizing a fluorescently labeled, biotinylated (dT)_45_ oligonucleotide to their poly(A) tails (Figure 1B) (Çetin et al., 2020). We then delivered Cy5-labeled eIF4E (O’Leary et al., 2013) (Supplemental Figure 1B,C) to the mRNAs.

eIF4E delivery led to transient cycles of mRNA–eIF4E single-molecule FRET (Figure 1C), as observed previously (Çetin et al., 2020; O’Leary et al., 2013). FRET was observed uniformly for the present set of 0.4 – 2 kb mRNAs, suggesting eIF4E is within ∼7 nm of the poly(A) tail when bound to the cap. We previously showed that almost all eIF4E–mRNA binding events result in FRET. However, to test the possibility that eIF4E induced mRNA end-to-end proximity, we Cy5-labeled the 5ʹ triphosphate groups of polyadenylated *NCE102* and *HXT2* (∼1.1 kb and ∼1.8 kb) (Lai et al., 2018), then immobilized them by poly(A) capture. (Supplemental Figure 2A). We observed persistent FRET between mRNA 5ʹ ends and poly(A) tails (Supplemental Figure 2B,C) for the majority of the Cy5 lifetime (Supplemental Figure 2D). These results are consistent with mRNA folding and tertiary compaction bringing the 5ʹ and 3ʹ ends within FRET distance (Lai et al., 2018; Khong and Parker, 2020; Vicens et al., 2018; Yoffe et al., 2011). Thus, our smFRET signal reliably reports on eIF4E–mRNA interaction (Supplemental Discussion D1).

We determined eIF4E–mRNA binding and dissociation rates by exponential fitting of the distributions of waiting times between binding events, and of the event durations (Çetin et al., 2020). Interactions differed substantially: association rates ranged over about four-fold, from 6.3 ± 1 µM^−1^ s^−1^ for *JJJ1* to 23 ± 2 µM^−1^ s^−1^ for *NCE102* (Figure 1D); dissociation rates depended less on mRNA identity (0.42 ± 0.05 s^−1^ to 0.9 ± 0.06 s^−1^) (Figure 1E; Supplemental Table 2). Thus, bimolecular eIF4E–mRNA affinity is more sensitive to how rapidly eIF4E associates with the cap than to how long it remains bound.

Assuming a two-state, on-off equilibrium-binding model (*K*_d_ = *k*_off_ / *k*_on_), eIF4E–mRNA affinities spanned a *K*_d_ range of ∼18 ± 6 nM to ∼96 ± 58 nM (Figure 1F), a higher affinity than for the dinucleotide cap analog, and equal or greater affinity than for capped oligonucleotides (Carberry et al., 1989; O’Leary et al., 2013; Ueda et al., 1991). However, whilst high-affinity, the bimolecular eIF4E–mRNA interaction is highly dynamic on the initiation timescale.

### eIF4E binds longer mRNAs more slowly

eIF4E–mRNA binding *in vivo* and translation dependence on eIF4G anticorrelate with mRNA and coding-sequence length (Costello et al., 2015; Park et al., 2011; Sen *et al*., 2016). This has been interpreted in terms of more efficient formation of the “closed-loop” mRNP, through eIF4G– poly(A) binding protein interaction, on shorter mRNAs (Thompson and Gilbert, 2017). However, it remained unclear whether eIF4E– and or eIF4E•G–mRNA dynamics might also vary with mRNA length.

We found no correlation of eIF4E association rates with leader length (Supplemental Figure 1D). However, mRNAs with longer coding sequences bound eIF4E more slowly (Figure 1G) (Pearson’s *R* = –0.786, *p* = 0.02). As the CDS accounts for 48% – 90% of the total length for these mRNAs, a similar correlation holds for total mRNA length (Supplemental Figure 1E). Occluded-volume effects on eIF4E diffusion in zero-mode waveguides due to increased mRNA size are highly unlikely decelerate eIF4E association, since the hydrodynamic radii of mRNAs in the 0.7 – 2.0 kb length range (7 – 12 nm; Borodavka et al., 2016) are at most ∼8% of the zero-mode waveguide diameter (∼150 nm; Moran-Mirabal and Craighead, 2008; Chen et al. 2014) Thus, eIF4E–mRNA association rates differ by over five-fold across a CDS length range that encompasses transcripts between the 10^th^ and 70^th^ percentiles of length in the yeast transcriptome (Tuller et al. 2009). This raised the question of whether similar steric effects would operate for eIF4E•eIF4G or eIF4F.

### Yeast eIF4G1 accelerates eIF4E–cap binding to differing extents for different mRNAs

eIF4G (Supplemental Figure 1B) accelerated eIF4E-mRNA association for all mRNAs (Figure 2A). However, the extent of acceleration differed by mRNA, from ∼7-fold for *JJJ1* to ∼4-fold for *NCE102*, resulting in rates from 42 ± 7 µM^−1^ s^−1^ to 84 ± 14 µM^−1^ s^−1^ (Supplemental Table 3). Acceleration also correlated significantly with CDS and total mRNA lengths (Figure 2B; Supplemental Figure 3A); shorter mRNAs showed less acceleration. Nevertheless, eIF4E binding remained faster for shorter mRNAs, though the range of rates is narrower than for eIF4E alone.

**Figure 2.**
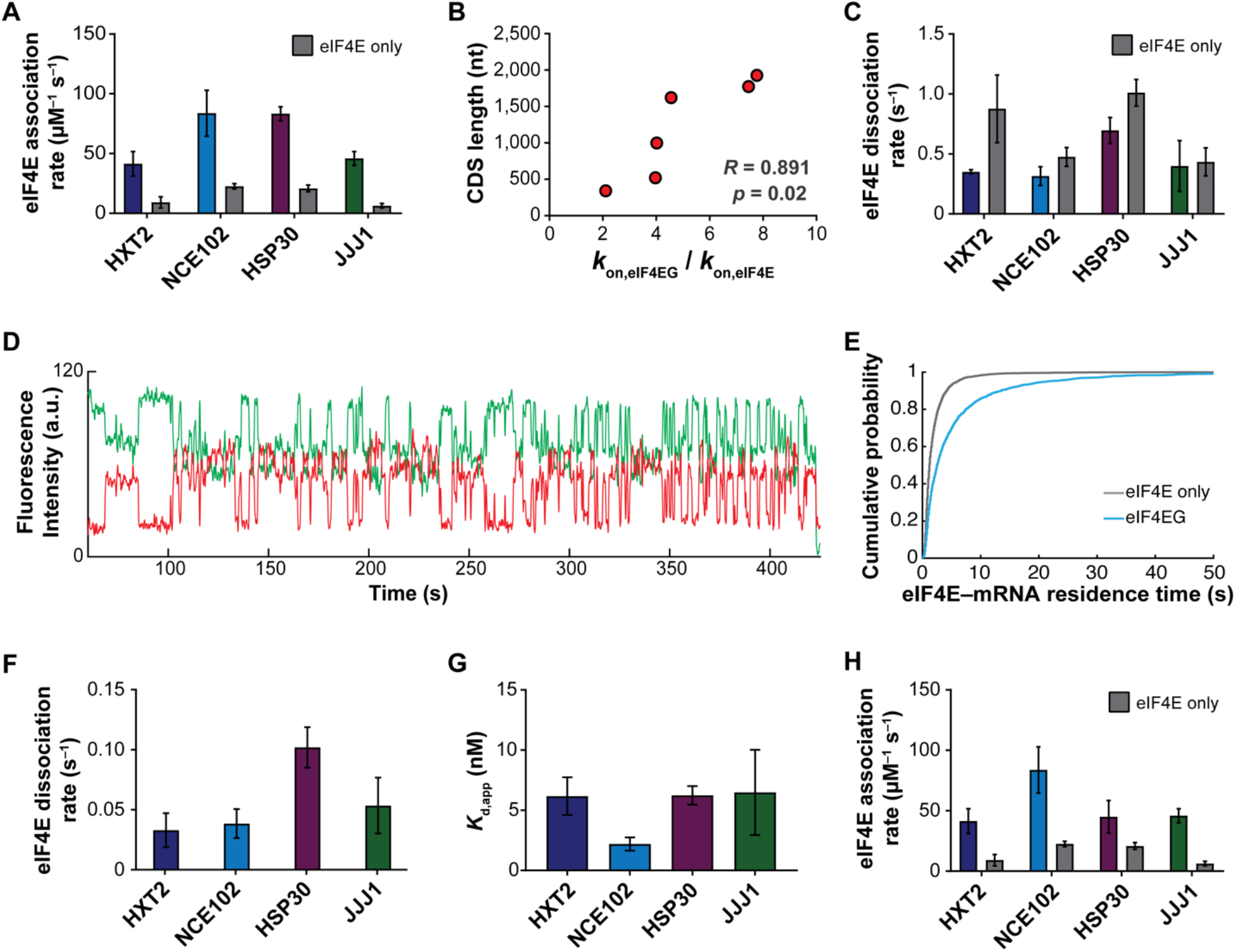
eIF4G1 accelerates eIF4E–mRNA binding in an mRNA-dependent manner and allows the interaction to persist on the translation initiation timescale. **A**. eIF4E-mRNA association rates in the presence of full-length yeast eIF4G1, compared with eIF4E-only rates. **B.** Fold-stimulation of eIF4E–mRNA association rate by eIF4G1, as a function of mRNA length. **C.** Kinetics of eIF4E–mRNA dissociation for transient binding events in the presence of eIF4G1. **D.** Representative single-molecule fluorescence trace for eIF4E–mRNA binding in the presence of eIF4G1. **E.** Cumulative probability distributions of eIF4E–*NCE102* mRNA event durations in the absence (grey) and presence (blue) of eIF4G1, showing appearance of slowly-dissociating events when eIF4G1 is present. **F**. eIF4E–mRNA dissociation rates for long-lived binding events in the presence of eIF4G1. **G.** Apparent equilibrium dissociation constants for the eIF4E–mRNA interaction in the presence of eIF4G1. **H.** eIF4E–mRNA association rates in the presence of eIF4G1_1–452_.

The narrowed association-rate range is consistent with cap accessibility being more similar between mRNAs in the presence of eIF4G. Furthermore, an eIF4G1 fragment (eIF4G_1–452_) containing the N-terminal RNA-binding domain but truncated immediately N-terminal to the eIF4E-binding domain, showed comparably accelerated eIF4E association (Figure 2H), consistent with eIF4G–mRNA binding playing a deterministic role in this enhanced cap accessibility.

More frequent eIF4E binding in the presence of eIF4G raised the possibility that the additional events were due to eIF4E•eIF4G binding to internal mRNA regions without eIF4E binding the cap. Experiments with uncapped (5ʹ-triphosphate), polyadenylated *NCE102* mRNA did result in transient eIF4E–mRNA FRET events (mean duration ∼1 ± 0.31 s), but at only ∼2% of the association rate relative to the capped mRNA (Supplemental Figure 3B–D). This is consistent with residual eIF4E•eIF4G binding either at the 5ʹ end, or at the 3ʹ end close to the FRET donor, and bolsters the conclusion that our FRET signal is specific for cap binding.

This result suggests that, while eIF4E•eIF4G–mRNA association is dominated by eIF4G, the cap structure offers a point of attachment that biases stable accommodation of eIF4E•eIF4G at the 5ʹ end. The results are also consistent with previous results of mammalian eIF4F binding to RNAs (Kaye *et al*., 2009), in that the affinity of eIF4F for RNA is driven dominantly by eIF4G.

### Yeast eIF4G1 allows the eIF4E–cap interaction to persist on the initiation timescale

Our data were surprising in that cap-binding events were short relative to the initiation timescale (Acker et al. 2009; Palmiter 1975), contrasting with eIF4E remaining associated with the 48S ribosomal pre-initiation complex throughout scanning (Bohlen et al., 2020). Fast eIF4E dissociation would limit how long an eIF4F•mRNA complex is available for PIC recruitment. On the other hand, different models have been proposed for whether eIF4E remains cap-bound as the mRNA 5ʹ end enters the PIC mRNA channel (Archer et al., 2016; Shirokikh and Preiss, 2018). eIF4E remaining cap-bound, or dissociating and rapidly rebinding, would favor maintaining eIF4G bound to the mRNA 5’ end, and thus looping of the leader as it moves through the PIC in search of the start site (Querido et al., 2020; Archer et al., 2016; Marintchev et al., 2009; Paek et al., 2015) We therefore assessed how eIF4G impacted the eIF4E–cap binding duration.

eIF4G slightly or moderately lengthened transient eIF4E–mRNA binding events (*k*_off_ = 0.30 ± 0.01 s^−1^ to 0.70 ± 0.08 s^−1^) (Figure 2C; Supplemental Table 3), resembling its effect on eIF4E binding to capped oligonucleotides (O’Leary et al., 2013). The effect varied between mRNAs, yielding a slightly narrower range of dissociation rates. However, the interactions remained transient on the initiation timescale.

Strikingly, though, a proportion of eIF4E-mRNA binding events lengthened by an order of magnitude in the presence of eIF4G (∼17% – 27%, depending on mRNA) (Figure 2D; Supplemental Table 3). The effect was observed as double-exponential behavior in the event-duration distribution (Figure 2E), and varied between mRNAs (*k*_off_ = 0.03 ± 0.01s^−1^ to 0.10 ± 0.01 s^−1^) (Figure 2F). These long events were not observed with the eIF4G_1–452_ fragment (Supplemental Figure 3E,F), implying they result from direct eIF4E–eIF4G interaction.

These rates exceed that for eIF4E dissociation from the eIF4G eIF4E-binding domain (*k*_diss_ ∼0.01 s^−1^ (Gross et al., 2003), which would suggest that eIF4E may dissociate from the mRNA while eIF4G remains bound. However, we cannot fully exclude that the events may be terminated by eIF4E•eIF4G dissociating from the mRNA as a unit. Indeed, we observed both behaviors for the eIF4F•mRNA complex in multicolor single-molecule fluorescence experiments described below. As the Cy5 lifetime is greater than two minutes in our illumination conditions (Supplemental Figure 2B,C), disappearance of FRET is unlikely to be due to Cy5 photobleaching.

The combined effects of eIF4G on association and dissociation drastically enhanced the apparent eIF4E–cap affinity: *K*_d,app_ values for the interaction were reduced ∼4.8 – 15.8-fold and became more similar between mRNAs (*K*_d_ range: 2.8 ± 0.9 nM to 7.3 ± 1.9 nM) (Figure 2G). Taken together, these data point to the eIF4G RNA-binding activity as central to both accelerating cap recognition and allowing it persist on the initiation timescale. While the accelerative effect is more pronounced on longer mRNAs, shorter mRNAs still bind eIF4E•eIF4G faster. This suggests a potential rationale for the finding that depleting eIF4G *in vivo* reduces translation globally, but compresses the range of translation efficiencies (Park *et al*., 2011) – in this rationale, shorter mRNAs sustain sufficient eIF4E•eIF4G–mRNA binding rates at lower eIF4G concentrations, but longer mRNAs become comparatively disadvantaged. Other factors that interact with eIF4F, such as Ded1p, could further modulate the length of this interaction.

### Free yeast eIF4A accelerates eIF4E-cap association in an mRNA-dependent manner

Free eIF4A (i.e., not eIF4F-bound) is present at ∼20 µM *in vivo* (von der Haar and McCarthy, 2002; Ghaemmagami *et al*., 2003; Lu *et al*., 2007), in large excess over eIF4E•eIF4G. Despite its abundance, cellular eIF4A appears to be rate-limiting for translation – i.e., small reductions in concentration substantially reduce translational output (Firczuk et al., 2013). We recently reported that free yeast eIF4A increased eIF4E association rates across mRNA populations (Çetin et al., 2020). However, it remained to test whether mRNAs are affected equally.

eIF4A and ATP addition indeed accelerated eIF4E-cap association for each mRNA (Figure 3A,B). However, the fold-acceleration ranged from 1.3 ± 0.1 (marginal, if any, stimulation) to 4.5 ± 0.8-fold, yielding rates from 24.8 ± 0.1 µM^−1^ s^−1^ to 52.8 ± 7.2 µM^−1^ s^−1^ (Supplemental Table 4). However, acceleration was quantitatively indistinguishable when ATP was substituted by the slowly-hydrolyzable analog ATP-*y*-S (Figure 3C; Supplemental Table 5). We performed the ATP-*y*-S experiments with the *JJJ1* and *NCE102* mRNAs as they showed substantial eIF4A enhancement of eIF4E binding. We opted for ATP-*y*-S, rather than the non-hydrolyzable AMPPNP, because AMPPNP does not support PIC–mRNA recruitment (Yourik et al., 2017). Nucleotide binding to eIF4A is thus sufficient for the rate acceleration. Meanwhile, eIF4E–mRNA binding events with free eIF4A and ATP remained essentially the same length relative to the eIF4E-only condition (Figure 3D). Taken together, these data are consistent with a model where eIF4A clamps the mRNA conformation in an ATP-dependent manner (Linder and Jankowsky, 2011), rendering the cap more accessible to eIF4E. The data are also consistent with proposed eIF4A roles in preventing RNA-RNA interactions that form polymeric condensates *in vitro* and stress granules and P-bodies *in vivo*, and in facilitating RNA-binding protein interactions (Linder and Jankowsky, 2011; Tauber et al., 2020).

**Figure 3.**
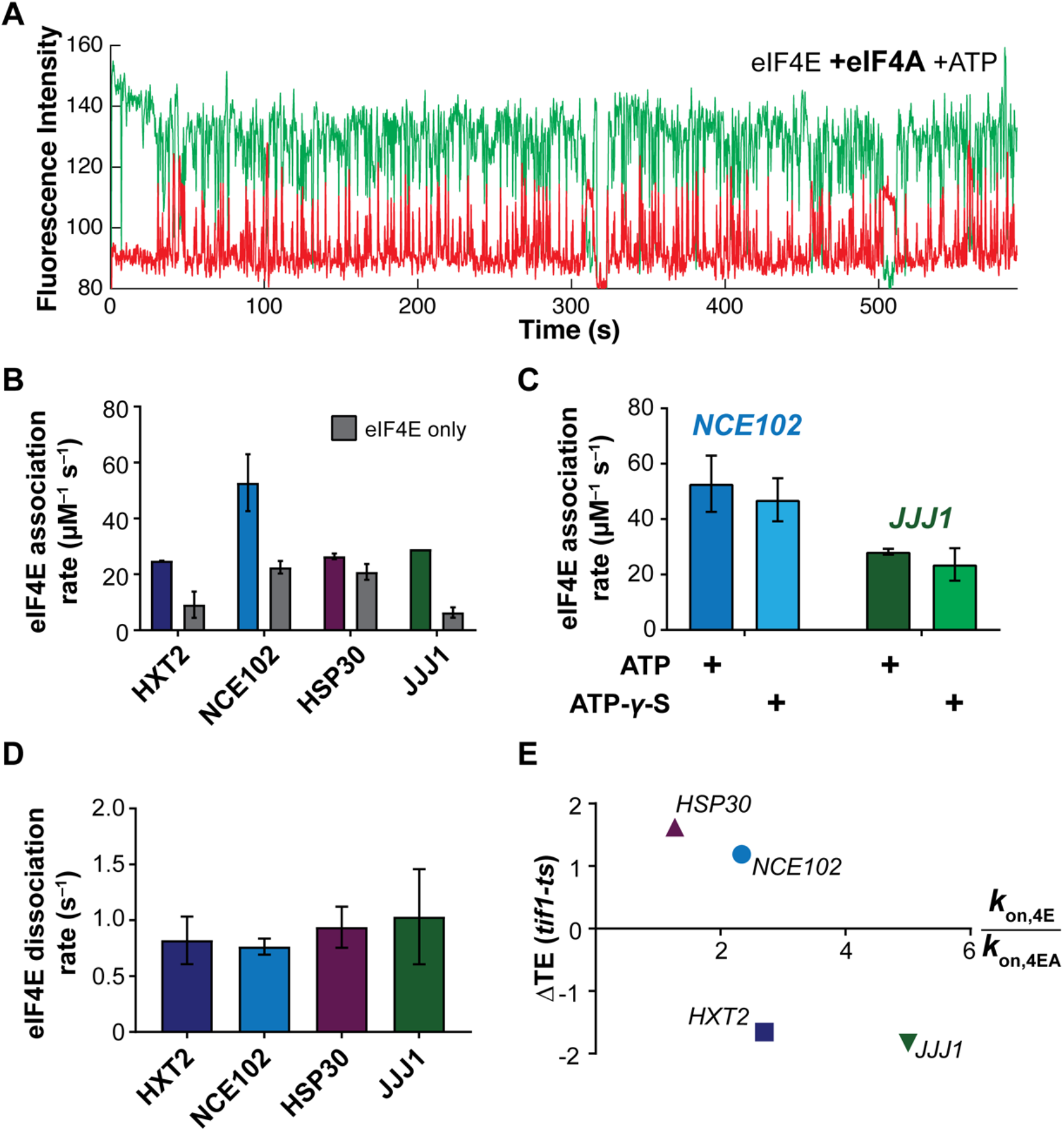
Free eIF4A with bound ATP stimulates eIF4E–mRNA association independently of eIF4G. **A.** Representative single-molecule trace showing eIF4E–mRNA interaction in the presence of 2 µM eIF4A and 2.5 mM ATP. **B.** eIF4E-mRNA association rates in the presence of eIF4A and ATP, compared with eIF4E-only rates. **C.** eIF4E–mRNA association rates in the presence of eIF4A and ATP or ATP-γ-S, for the *NCE102* and *JJJ1* mRNAs. **D.** eIF4E–mRNA dissociation rates in the presence of free eIF4A and ATP. **E.** Relationship between translation-efficiency dependence on eIF4A and the fold-increase in eIF4E–mRNA association rate induced by free eIF4A and ATP.

As observed for eIF4G, the fold-acceleration of eIF4E–mRNA binding induced by eIF4A again was greater for longer (*JJJ1*, *HXT2*) *vs.* shorter (*NCE102*, *HSP30*) mRNAs. This would be consistent with modulation of tertiary structure by eIF4A–RNA binding as a mechanism for the rate enhancement. In parallel, mRNA-to-mRNA differences for this eIF4A activity could stem from mRNA features that lead to different extents of mRNA single- or double-strandedness, for which yeast eIF4A has significantly different affinities (Rajagopal *et al*., 2012).

To probe the relevance of this stimulation to translation *in vivo*, we correlated our kinetics with ribosome-profiling data for the mRNAs’ eIF4A–translation efficiency dependence (Sen et al., 2015). eIF4A indeed accelerates eIF4E binding to a greater degree for mRNAs that are hyperdependent (*HXT2, JJJ1*) on eIF4A compared to hypodependent mRNAs (*HSP30, NCE102*) (∼1.8 vs. 3.9-fold) (Figure 3E). Thus, free eIF4A may contribute significantly to translation efficiency right from the outset of initiation, by promoting cap accessibility for eIF4E binding and ribosome recruitment.

### eIF4F differentiates eIF4E–mRNA interactions in an ATP-dependent manner

We next assessed how the full eIF4F complex modulated eIF4E–mRNA binding. We included eIF4A (2 µM) in excess of eIF4G (250 nM) and eIF4E (10 – 30 nM) to limit nonspecific interaction of eIF4E with the ZMW waveguides, which interferes with data analysis at concentrations greater than this.

In the eIF4F complex without ATP, eIF4E–mRNA association accelerated on all mRNAs relative to eIF4E alone (Figure 4A,C), with rates between 43.1 ± 13.3 µM^−1^ s^−1^ for *JJJ1* and 104.8 ± 3.1 µM^−1^ s^−1^ for *HSP30* (Supplemental Table 6). This rate for *HSP30* was the fastest measured in the present study. Acceleration again varied between mRNAs, from 6.8 ± 1.9 to 3.0 ± 0.4-fold. The net effect was to differentiate eIF4E–mRNA binding between mRNAs relative to the eIF4E•eIF4G condition. Long and short eIF4E–mRNA binding events were also observed with eIF4F, and their relative incidence was unchanged within experimental error. Transient eIF4E dissociation occurred at around ∼0.4 s^−1^ – 0.6 s^−1^, similar to eIF4E•eIF4G (Figure 4D), while long events dissociated at ∼0.05 s^−1^ and 0.10 s^−1^ (Figure 4E), slightly faster than for eIF4E•eIF4G. These results again place eIF4G as a dominant kinetic contributor to eIF4F–mRNA affinity, echoing thermodynamic data for human eIF4F (Kaye et al., 2009).

**Figure 4.**
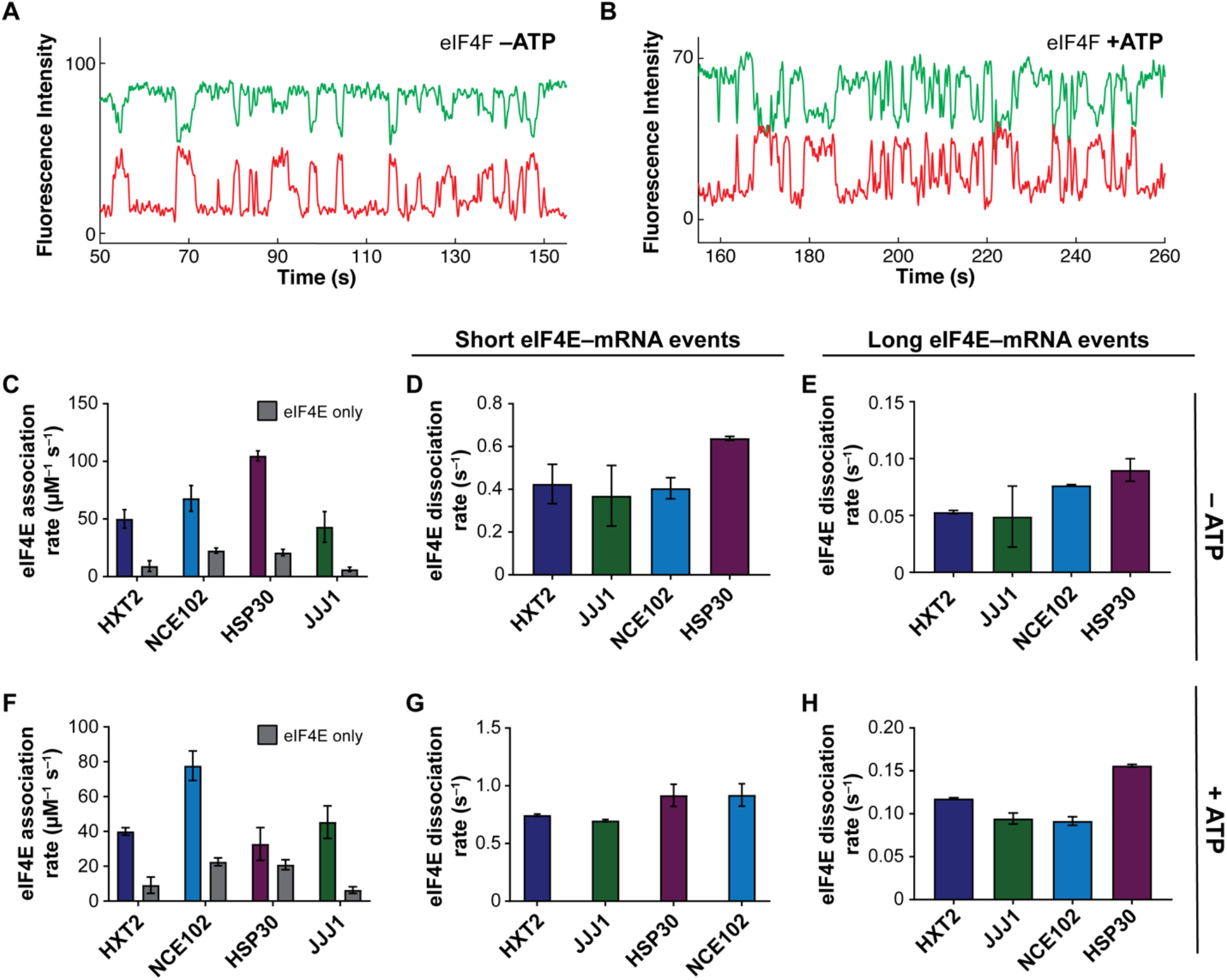
The eIF4F complex discriminates eIF4E–mRNA interaction dynamics in an ATP-dependent manner. **A.** Representative single-molecule fluorescence trace for eIF4E–mRNA interaction in the eIF4F complex without added ATP, on *NCE102*. **B** Representative trace for eIF4E–mRNA binding in the eIF4F complex with ATP, on *NCE102*. **C.** eIF4E–mRNA association rates in the eIF4F complex without ATP. **D.** Dissociation rates of transient eIF4E–mRNA interactions in the eIF4F complex without ATP. **E.** Dissociation rates of long-lived eIF4E–mRNA interactions in the eIF4F complex without ATP. **F.** eIF4E–mRNA association rates in the eIF4F complex with ATP. **G.** Dissociation rates of transient eIF4E–mRNA interactions in the eIF4F complex with ATP. **H.** Dissociation rates of long-lived eIF4E–mRNA interactions in the eIF4F complex with ATP.

However, addition of ATP led to both mRNA-specific and global changes in dynamics (Figure 4B,F–H). The *HSP30* association rate was strikingly reduced, from being the fastest among the mRNAs, to being the slowest (25.8 ± 5.0 µM^-1^ s^−1^) with ATP present (Figure 4F; Supplemental Table 7). This almost entirely reversed the acceleration in eIF4E–mRNA binding afforded to *HSP30* by eIF4F. On the other hand, the *NCE102*, *HXT2* and *JJJ1* mRNAs showed small or no reductions in association rate on ATP addition.

ATP addition also shortened both the long and short eIF4E binding events, i.e. the complex became more dynamic (Figure 4B,G,H). For the long events, this effect ranged from a modest ∼50% for *HSP30* to more than threefold for *HXT2*. Dissociation kinetics became even more similar between mRNAs than in the other conditions, pointing to a common rate-limiting step for eIF4E dissociation from the eIF4F•mRNA complex. Thus, coordinated activities of all eIF4F subunits determine the efficiency and duration of eIF4E–cap binding in eIF4F.

### Simultaneous direct observation of eIF4A– and eIF4E–mRNA interaction

To broaden our view of eIF4F intersubunit coordination during cap recognition, we performed three-color experiments that included fluorescent Cy3-eIF4A (Supplemental Figure 4A) with Cy5-eIF4E and Cy3.5-mRNA (Figure 5A). Cy3-eIF4A RNA-dependent ATPase activity was indistinguishable from the unlabeled protein (Supplemental Figure 4B,C). For these experiments we chose *JJJ1*, *NCE102*, and *HXT2*, which span the range of stimulation of eIF4E binding by eIF4A.

**Figure 5.**
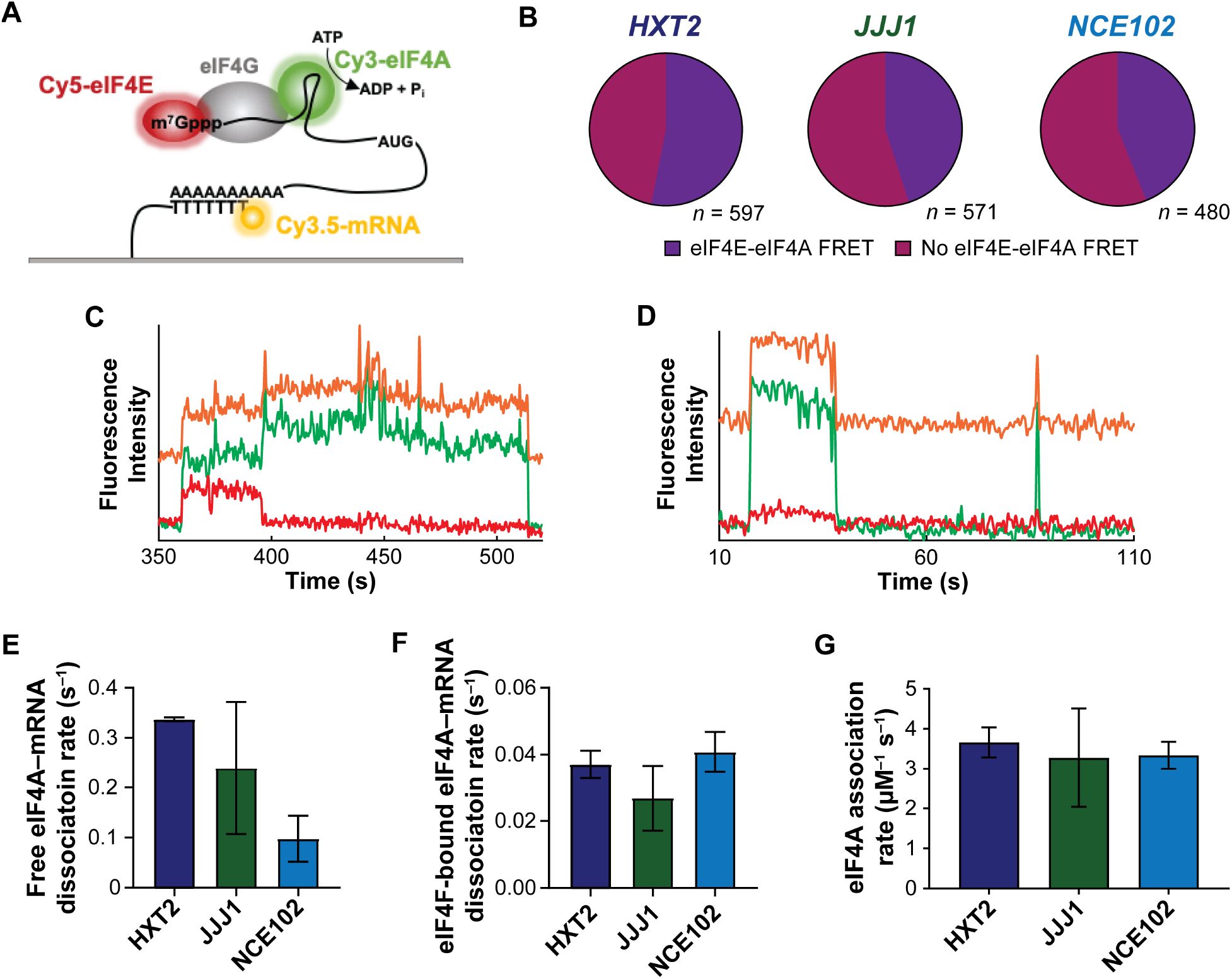
Three-color smFRET to probe eIF4F- and eIF4A-mRNA interaction dynamics. **A.** Schematic of the three-color smFRET experiment with two donors (on eIF4A and mRNA) and one acceptor (eIF4E). A FRET signal between Cy5-labeled eIF4E and the Cy3.5-labeled mRNA is tracked at the same time as a FRET signal between Cy3-eIF4A and Cy5-eIF4E. **B.** Relative incidence of eIF4A-mRNA binding occurring with and without FRET to eIF4E. *n* is the number of molecules analyzed to enumerate the event types on each mRNA. **C.** Representative smFRET trace showing concomitant mRNA binding of eIF4E and eIF4A with eIF4E–eIF4A FRET, consistent with eIF4F–mRNA binding, on *JJJ1*. **D.** Representative single-molecule fluorescence trace for free eIF4A–mRNA binding on *JJJ1*. The Cy3 and Cy5 signals were manually corrected by linear subtraction to equalize their background values, for clarity of presentation. **E.** eIF4A– mRNA dissociation rates following eIF4A–mRNA binding without eIF4E. The Cy3 and Cy5 signals were manually corrected by linear subtraction to equalize their background values, for clarity of presentation. **F.** eIF4A–mRNA dissociation rates following eIF4A–mRNA binding with eIF4E. **H.** eIF4A–mRNA association rates across all binding event types.

We observed two types of eIF4A–mRNA binding event. In the first, eIF4A binding was accompanied by eIF4E–eIF4A FRET. (43%, 53%, and 45% of all eIF4A binding events for *NCE102*, *HXT2*, and *JJJ1*, respectively) (Figure 5B,C). Since no direct eIF4E–eIF4A interaction is known, we interpret these events to represent assembly of an intact eIF4F complex. In a second event type, eIF4A bound mRNA without FRET to eIF4E (57%, 47%, and 55% for *NCE102*, *HXT2*, and *JJJ1*) (Figure 5D). Approximately 10% of eIF4A is expected to be free – i.e., not bound to eIF4E•eIF4G – under our conditions (Mitchell et al., 2010), consistent with observation of these events.

The dynamics of eIF4A–RNA binding events lacking eIF4E–eIF4A FRET varied between mRNAs and also differed kinetically from events where eIF4E–eIF4A FRET was observed. Two types of this binding mode were observed – transient and longer-lived, with the transient events constituting 25 – 90% of the eIF4A–mRNA encounters, depending on mRNA (Supplemental Figure 5; Supplemental Table 8). The dissociation rates for the dominant (higher-amplitude) eIF4A dissociation pathway varied considerably between the mRNAs, from 0.097 ± 0.032 s^−1^ (*NCE102*) to 0.337 ± 0.003 s^−1^ (*HXT2*) (Figure 5E; Supplemental Table 8). Conversely, the eIF4A–mRNA dissociation rates in eIF4A–eIF4E co-binding events were kinetically more similar between mRNAs, and ranged from 0.0269 s^−1^ ±0.004 (*JJJ1*) to 0.04077±0.003 (*HXT2*) (Figure 5F; Supplemental Table 9).

eIF4A–mRNA association rates were identical between mRNAs within experimental error (Figure 5G, Supplemental Table 8). Interestingly, then, and in contrast to eIF4E, variable affinity of free eIF4A for different mRNAs appears to result from differences in the lifetimes of the eIF4A•mRNA complexes. This echoes results that demonstrate different conformational dynamics of eIF4A in the presence of RNA oligonucleotides that differ in their duplex properties (Rajagopal *et al*., 2012; Andreou et al. 2019). Extrapolating our data to cellular concentrations of eIF4A, our results implicate free eIF4A as a multifunctional “mRNA chaperone” that maintains cap accessibility for eIF4F binding.

### eIF4E is ejected from the cap after initial eIF4F–mRNA binding

A key question around cap recognition is how the mRNA 5ʹ end is transferred into its channel on the 40S subunit if the cap is bound by eIF4E/eIF4F. We found that eIF4E fluorescence frequently disappeared before eIF4A fluorescence after formation of an eIF4F•mRNA complex (Figure 6A). This was the most common outcome, and occurred for ∼51% of eIF4F–mRNA binding events on *HXT2*, 45% on *JJJ1*, and 56% on *NCE102* (Figure 6B). The next most common outcome was simultaneous disappearance of eIF4E and eIF4A fluorescence (33%, 43%, 29%). More rarely, eIF4E bound mRNA after eIF4A and then dissociated prior to eIF4A (16,12% 15%). Thus, there is a preference for disrupting eIF4E–cap interaction whilst maintaining eIF4A–mRNA binding. This echoes findings for mammalian eIF4F where cap binding appears to reduce eIF4E affinity for eIF4F (Merrick, 2015; Ray et al. 1985).

**Figure 6.**
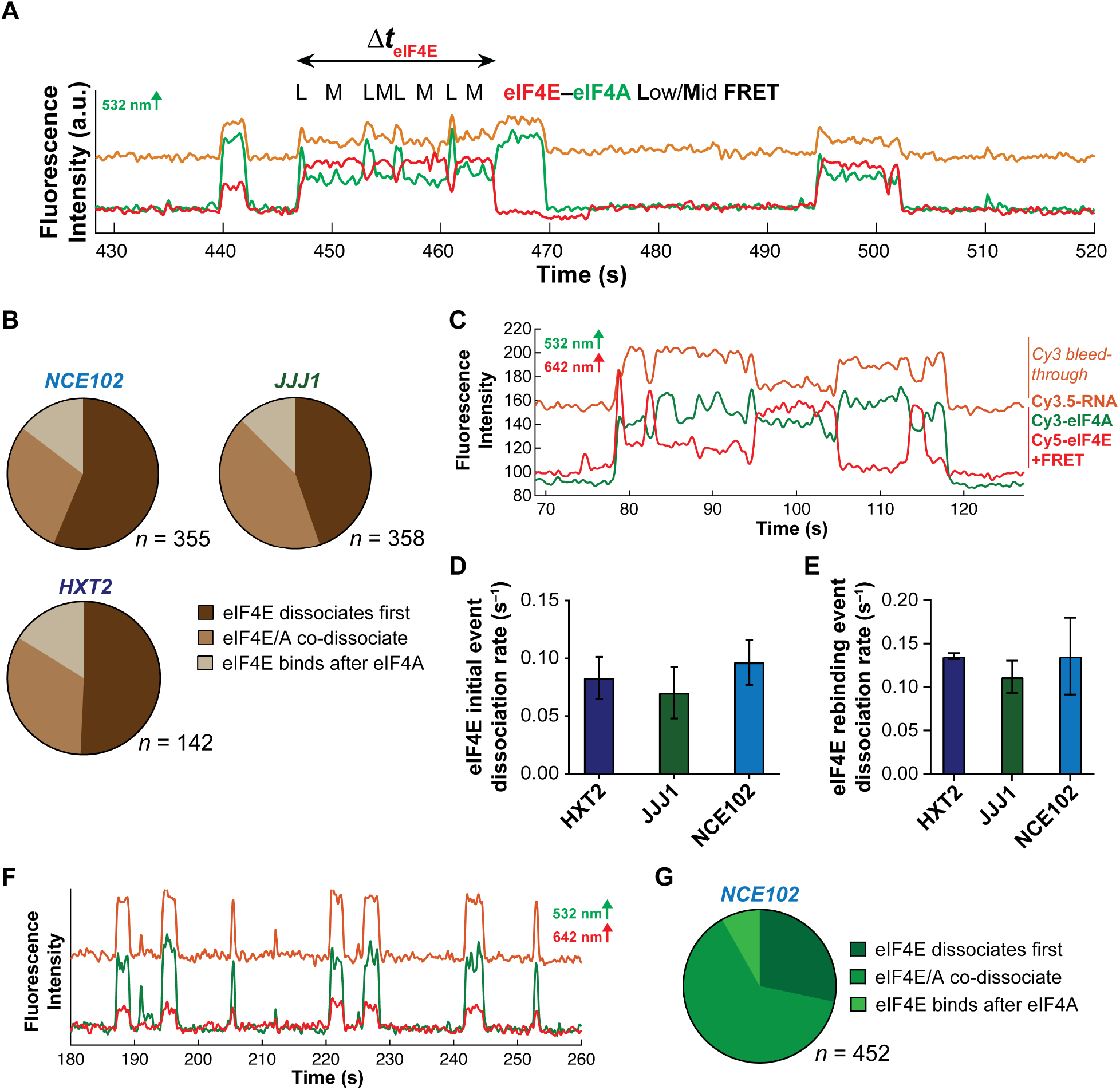
Dynamic coordination within eIF4F after cap recognition. **A.** Representative smFRET trajectory showing event with ejection of eIF4E prior to eIF4A, and fluctuations in the eIF4F•mRNA conformation on *JJJ1*. **B.** Relative incidence of eIF4E or eIF4A dissociation, or co-dissociation from eIF4F•mRNA complexes. *n* is the number of molecules analyzed to enumerate the event types on each mRNA. **C.** Representative single-molecule fluorescence trajectory for eIF4F•mRNA complex formation and dynamics, observed by dual red/green illumination. **D.** Rates for the initial eIF4E–mRNA dissociation event after eIF4F–mRNA complex formation. **E.** eIF4E– mRNA dissociation rates for events where eIF4E rebinds mRNA following initial dissociation from eIF4F•mRNA. **F.** Representative single-molecule fluorescence trajectory for eIF4F•mRNA complex dynamics with ATP-γ-S. The Cy3 and Cy5 signals were manually corrected by linear subtraction to equalize their background values, for clarity of presentation. **G.** Relative incidence of eIF4E or eIF4A dissociation, or co-dissociation from the eIF4F•mRNA complex in the presence of ATP-γ-S.

Disappearance of eIF4E fluorescence could be due either to dissociation from eIF4F•mRNA, or adoption of an extended eIF4F conformation that places eIF4E out of FRET range to both mRNA and eIF4A. To differentiate between these possibilities, we repeated the experiment with direct excitation of the Cy5-eIF4E fluorophore. In this illumination scheme, all eIF4E–mRNA interactions are detected, rather than only interactions that produce FRET – i.e., adoption of a no-FRET conformation would be reported by loss of FRET but persistence of the Cy5 signal. We found that disappearance of eIF4E–mRNA FRET following eIF4F–mRNA binding was due to complete dissociation of Cy5-eIF4E from mRNA for 90% of the FRET events on the *JJJ1* mRNA (Figure 6C). Put otherwise, eIF4E is ejected from the eIF4F•mRNA complex shortly after cap recognition.

We also observed relatively frequent eIF4E rebinding after initial ejection (Figure 6C). However, while the first eIF4E dissociation event occurred at a rate of 0.07 – 0.08 s^−1^ across all mRNAs, dissociation during the subsequent rebinding events was slightly faster (*k*_off_ ∼0.10 s^−1^ – 0.13 s^−1^) (Figure 6D,E; Supplemental Table 10). Importantly, though, they were still not as fast as eIF4E dissociation in the absence of eIF4G (Figure 1E), further arguing that they are due to eIF4E rebinding with eIF4G still bound to the mRNA. Evidently a change occurs in the eIF4F complex once it has established itself at the mRNA 5ʹ end which enhances eIF4E dissociation. Indeed, a regular feature of the eIF4F•mRNA complex was fluctuation of the FRET efficiency during the binding events (Figure 6A), suggesting the occurrence of conformational rearrangements.

To establish the role of ATP hydrolysis in ejection of eIF4E from the eIF4F•mRNA complex, we substituted ATP with ATP-γ-S and monitored the dynamics of the corresponding eIF4F•mRNA complexes (Figure 6F). With the slowly-hydrolyzable analog, the relative incidence of eIF4E– eIF4A co-dissociation from RNA increased at the expense of eIF4E ejection (Figure 6G). Thus, ejection of eIF4E appears attributable to ATP hydrolysis in the eIF4F complex.

## DISCUSSION

### An mRNA-centric model for cap-recognition dynamics

Our results reveal that eIF4F interactions with full-length mRNAs are highly dynamic on the initiation timescale, and that the dynamics vary substantially between mRNAs. Variability is dominantly due to differences in the rate of factor–mRNA binding, which is influenced by mRNA features and is tuned by eIF4F subunits and the presence of ATP. Taken together, the data suggest a model where a topographically condensed mRNA molecule initially has limited 5ʹ cap accessibility for eIF4E binding (Figure 7). This steric block results in a relatively wide variation of eIF4E–mRNA association rates, and is relieved by RNA-binding activities of eIF4G and free eIF4A. For eIF4A, there is a distribution of labor between the free and eIF4F-bound enzyme. That the ability of free eIF4A to promote eIF4E–mRNA binding relates to translation dependence on eIF4A *in* vivo suggests an explanation for why, despite its high abundance, modest reductions in cellular eIF4A concentration reduce translational output (Firczuk *et al*., 2013). Our results are also consistent with the observation that elements throughout the mRNA are involved in acceleration of mRNA–PIC recruitment by eIF4A (Yourik et al., 2017). Moreover, they are reminiscent of roles proposed for eIF4A in modulating RNA condensation and stress-granule formation (Tauber et al., 2020). Additional RNA helicases (Gao et al., 2016), RNA-binding proteins targeted to specific mRNA features (Hentze et al., 2018), or indeed general RNA-binding proteins (Svitkin et al., 2009) are also predicted to impact eIF4E–mRNA dynamics in this model, opening additional avenues for broad or targeted regulation of cap recognition

**Figure 7.**
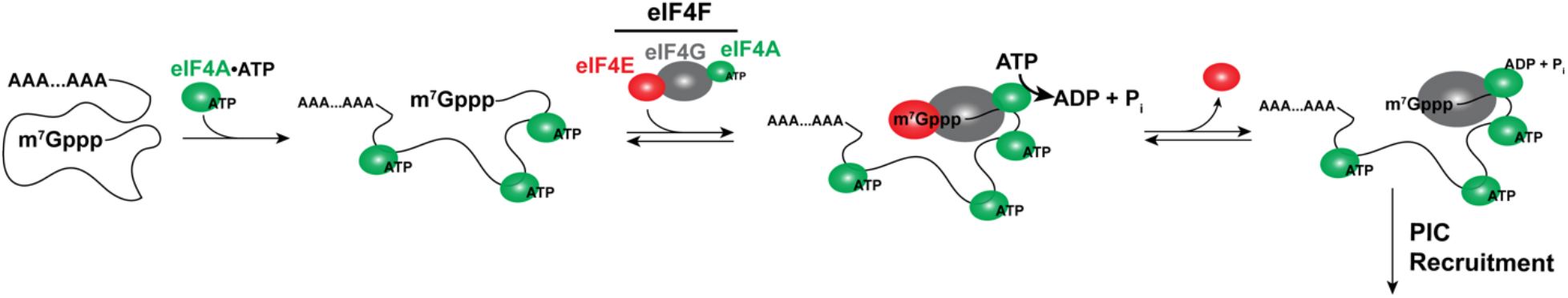
Mechanistic model for eIF4F–mRNA interaction dynamics.

### mRNA length sensing during early initiation

Our results suggest that eIF4E– and eIF4F–mRNA dynamics are sensitive to coding-sequence and mRNA length, and that RNA-binding activity of individual initiation factors mitigates this sensitivity. Because both eIF4G and eIF4A, which bind RNA through different structural mechanisms, can both accelerate eIF4E binding, our data point to disruption of tertiary contacts in longer mRNAs that sterically oppose eIF4E–cap attachment as the source of the length dependence. These findings augment, or perhaps invert, the previously-proposed model that closed loop formation is more efficient on short mRNAs (Thompson and Gilbert, 2017). Indeed, cryo-EM studies have found that polysomes remain circular even without mRNA polyadenylation (Afonina et al., 2014; Merrick, 2015), even in actively translating cell lysates. Instead of poly(A)-binding protein being more efficient at recruiting eIF4E•eIF4G on shorter mRNAs, our results argue that shorter mRNAs may support intrinsically faster eIF4E binding, allowing more rapid formation of eIF4G–Pab1p interactions.

Importantly though, our data do not exclude a role for poly(A)-binding protein in modulating eIF4F•mRNA dynamics. PABP–eIF4G binding could alter eIF4F conformational dynamics to increase the likelihood of efficient binding of eIF4E to the mRNA cap. This would be consistent with the ability of poly(A)-binding protein to stimulate translation on non-polyadenylated mRNAs when provided poly(A) *in trans* (Munroe and Jacobson, 1990). Moreover, several classes of mRNAs have been identified with differing relative enrichments in eIF4E•eIF4G and Pab1p (Costello *et al*., 2015), pointing to the potential for additional modes of crosstalk between these factors in cap recognition. Interestingly, this may differ in mammals, where the UTR regions comprise a greater portion of the mRNA length, and ORF length appears to have a reduced impact on translational efficiency (Riba et al., 2019).

### Dynamic eIF4F subunit coordination in the eIF4F•mRNA complex

Once eIF4F is bound to the mRNA, yeast eIF4G–mRNA binding maintains the eIF4E–cap interaction on the initiation timescale, and the complex is conformationally dynamic. Lack of structural information for the full eIF4F•mRNA complex renders assessing the nature of these conformational rearrangements challenging. However, a structure of the human 48S PIC that included partial density for eIF4G indicated that the protein is highly flexible; likewise, a structure of the yeast 48S PIC did not identify the position of eIF4F, attributed to its dynamic nature (Querido et al., 2020, Llácer et al. 2018), pointing to this as one potential source of the conformational rearrangements. Similarly, eIF4G stabilizes a closed conformation of eIF4A (Harms et al., 2014; Querido et al., 2020), and this is associated with a reciprocal conformational change in eIF4G (Schütz et al., 2008).

Our data suggest a molecular mechanism for guiding the mRNA as it accommodates onto the 40S subunit during PIC recruitment. Our FRET data indicate that the cap structure induces an intrinsic and strong kinetic bias toward eIF4F–5’ end binding through eIF4E–cap interaction. Once eIF4F is bound to mRNA, eIF4G maintains the eIF4E–cap interaction, but eIF4A-catalyzed ATP hydrolysis activity releases it, freeing the 5’ end whilst eIF4G and eIF4A remain mRNA-bound to act in contacting the PIC during its attachment to the mRNA (He *et al*., 2003; Yamamoto *et al.,* 2005). Since the PIC independently stimulates the eIF4A ATPase activity (Yourik *et al*., 2017), this suggests a model where initial PIC binding might gate release of eIF4E from the cap to allow mRNA–PIC accommodation. Transient eIF4E–cap rebinding events may then ensure that the complex is recycled and retained at the mRNA 5’ end for the next round of PIC recruitment after scanning, or for efficient reinitiation.

### Implications for translational control

Our results suggest a model for how eIF4E/eIF4F concentration can be leveraged to mediate differences in mRNA activation for translation, even though eIF4F–mRNA binding is saturated in an equilibrium sense under cellular conditions, where eIF4E and total mRNA are present at similar concentrations in yeast cells – ∼1 µM (Ghaemmagami *et al*., 2003; Lu *et al*., 2007) – in great excess over eIF4F•mRNA *K*_d_ values. Based on the eIF4E–mRNA binding rates we measured for the eIF4F complex, at these concentrations eIF4E will be substantially bound to mRNAs in cells, thus limiting the pool of eIF4F available for *de-novo* translation initiation (Pelletier *et al*., 2015; Chu *et al*., 2020). While we cannot exclude that the situation for polysomal mRNA may differ, the model is supported by the finding that small reductions in eIF4E concentration linearly reduce translational output (Firczuk et al., 2013).

In this regime, the rate of the eIF4E–mRNA interaction that begins *de-novo* cap recognition becomes an important determinant of initiation efficiency. Differences in this rate between mRNAs differentiate the mRNAs’ ability to compete for initiation, in accord with mathematical modeling (Godefroy-Colburn and Thach, 1981). Given our finding that mRNA length impacts the eIF4E association rate, these results highlight how information encoded along the mRNA length intrinsically contributes to the efficiency of early initiation, and offers one explanation for why longer mRNAs are often translated less efficiently and with higher eIF4F dependence (Thompson and Gilbert, 2017; Costello *et al.,* 2015; Sen *et al*., 2016). Moreover, reduction in the availability or activity of any one eIF4F subunit is expected to impact different mRNAs differently in this model. Short mRNAs and/or less structured mRNAs that effectively compete for eIF4F–cap binding are predicted to be less sensitive to depletion of active eIF4F. mRNAs that associate faster with eIF4F might be expected to have an advantage in terms of translational efficiency under conditions such as stress where eIF4F activity is downregulated, provided they do not accumulate in stress granules or P bodies.

A second aspect of this kinetic scheme is that eIF4G, by inducing long eIF4E–mRNA binding events, has the effect of rendering cap recognition a non-equilibrium process on the initiation timescale. PIC–mRNA recruitment rates measured *in vitro* are in the ∼0.001 – 0.01 s^−1^ range at 30 nM 40S subunits (Mitchell et al., 2010; Yourik et al., 2017). Extrapolating to a ∼1 µM lower limit for the cellular concentration of free 40S subunits thus implies a minimum recruitment rate of ∼0.05 – 0.4 s^−1^ at physiological PIC concentrations. This places recruitment occurring at rates close to or much faster than slow eIF4E–cap dissociation in the presence of eIF4G. With this kinetic balance, the overall efficiency of all steps from initial cap recognition to PIC–mRNA attachment is partially or fully limited by the eIF4E–mRNA association rate if PICs are sufficiently abundant. In contrast, for mRNAs that associate rapidly with eIF4E, PIC availability is predicted to limit initiation, suggesting a rationale for why translation of different subsets of mRNAs is affected differently by regulation directed at eIF4F and the PIC (Gebauer and Hentze, 2004).

Interestingly, mammalian eIF4F differs from yeast eIF4F in several aspects, particularly in terms of eIF4G1 architecture and eIF4A helicase activity. Moreover, mammalian mRNAs are generally longer and more structurally complex due to their higher GC content. Hence, it will be important to separately characterize these dynamics for mammalian eIF4F in future experiments. In summary, we find that an intricate interplay of mRNA identity with factor activities facilitates mRNA-to-mRNA discrimination in cap recognition. Within this, ATP binding and hydrolysis also play important and distinct roles. The dynamics of the eIF4F•mRNA complex suggest a mechanism for coordinating eIF4F–cap recognition with PIC recruitment, and for conferring mRNA sensitivity to distinct translational control pathways.

## AUTHOR CONTRIBUTIONS

S.O’L. conceived the project. B.Ç. and S.O’L. designed the experiments. B.Ç. performed the experiments. B.Ç. and S.O’L. analyzed the data and wrote the manuscript. Both authors approved the final manuscript version.

## ACKNOWLEDGEMENTS

We thank Jin Chen (UT Southwestern) and Gregor Blaha (UC Riverside) for insightful discussions on the manuscript. This work was supported by funding from the National Institutes of Health (awards GM138939 and GM139056 to S.O’L.), and from UC Riverside (Regents’ Faculty Fellowship award to S.O’L.).

## DECLARATION OF INTERESTS

The authors declare no competing interests.

## SUPPLEMENTAL FIGURE TITLES AND LEGENDS

**Supplemental Figure 1.**
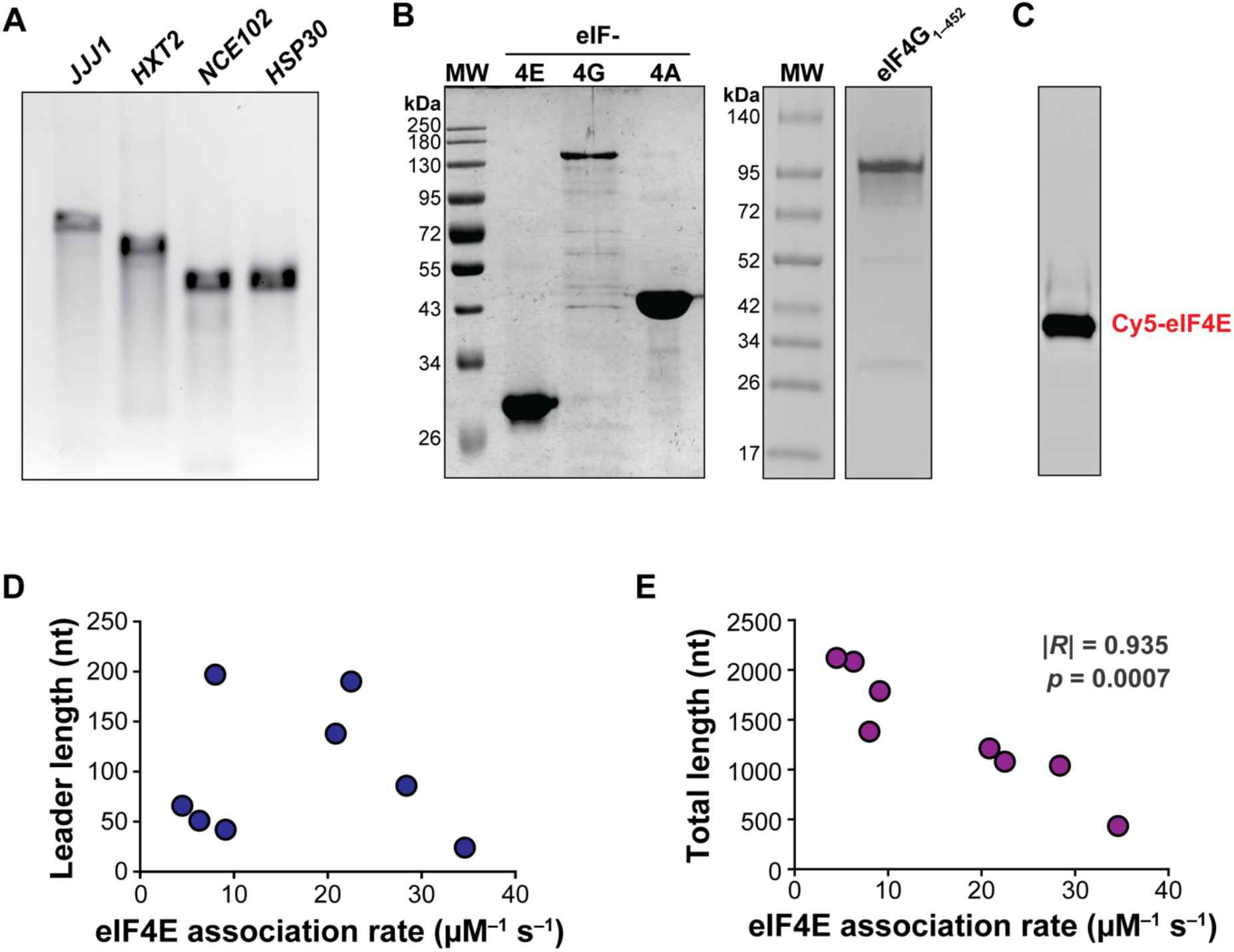
Relates to Figure 1. Preparation of mRNAs, recombinant proteins, and correlation of mRNA lengths to the eIF4E association rate. **A.** Representative agarose gel electrophoresis of mRNA transcripts after capping and polyadenylation. **B.** SDS-PAGE analysis of purified recombinant eIF4F factors. **C.** SDS-PAGE analysis of purified Cy5-eIF4E imaged by excitation of Cy5. **D.** Correlation of 5ʹ UTR lengths with eIF4E–mRNA association rates. **E.** Correlation of total mRNA lengths with eIF4E association rates. A significant correlation is observed between eIF4E binding rate and mRNA length.

**Supplemental Figure 2.**
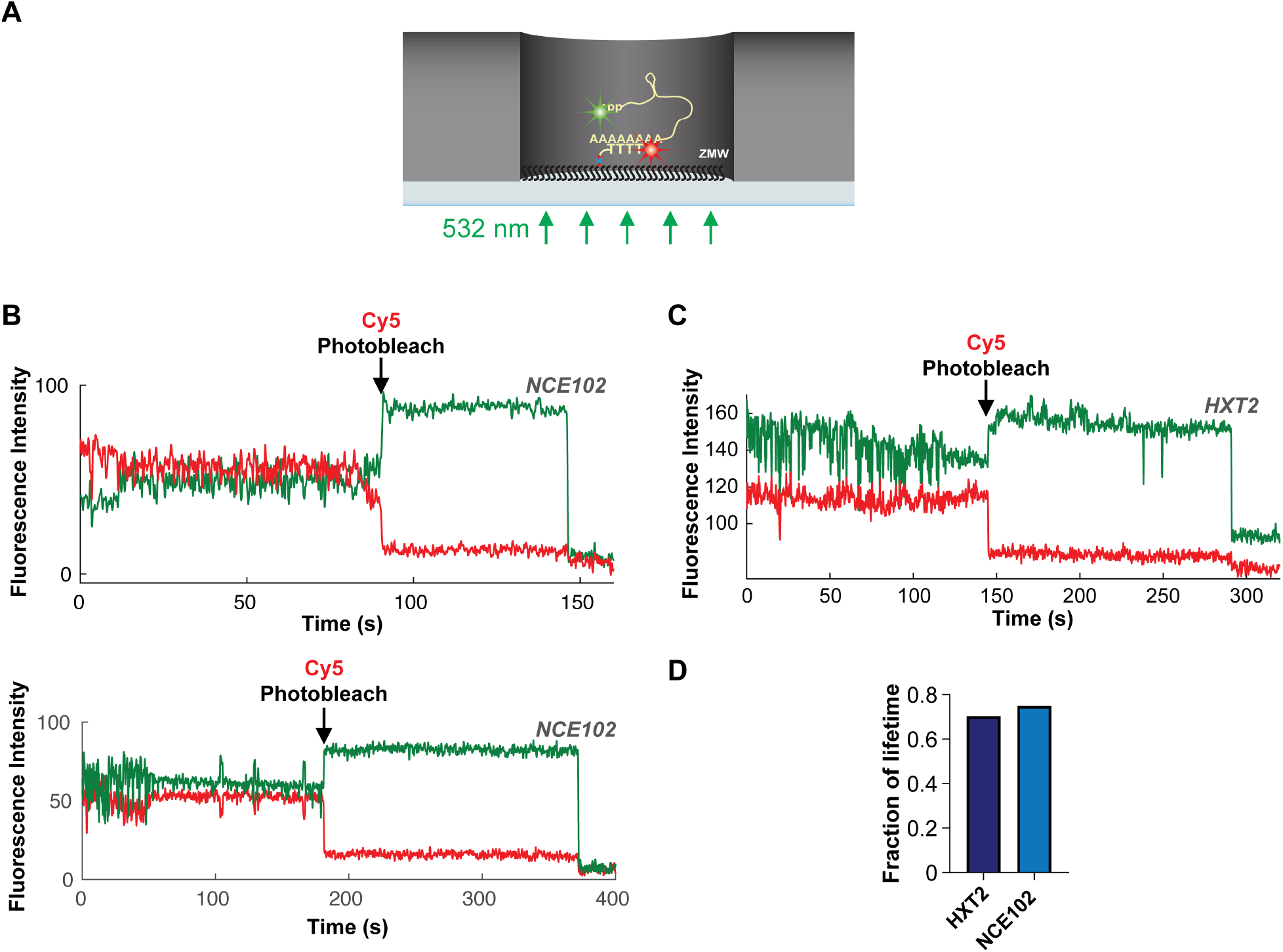
Relates to Figure 1. Intrinsic proximity of mRNA ends allows accurate determination of eIF4E–mRNA arrival rates. A. Schematic of intramolecular smFRET experiment to assess intrinsic mRNA end-to-end proximity. Uncapped mRNA is labeled through its 5ʹ triphosphate group with a FRET donor (Cy3), and on its poly(A) tail with a FRET acceptor (Cy5), then surface-immobilized through poly(A) capture. **B,C.** Representative smFRET traces showing 5’- to 3’-end FRET for two yeast mRNAs (*NCE102*, panel B; *HXT2, C*). Efficient smFRET is terminated by apparent acceptor photobleaching. **D.** Plot showing the fraction of Cy5 lifetime spent in a detectable FRET state.

**Supplemental Figure 3.**
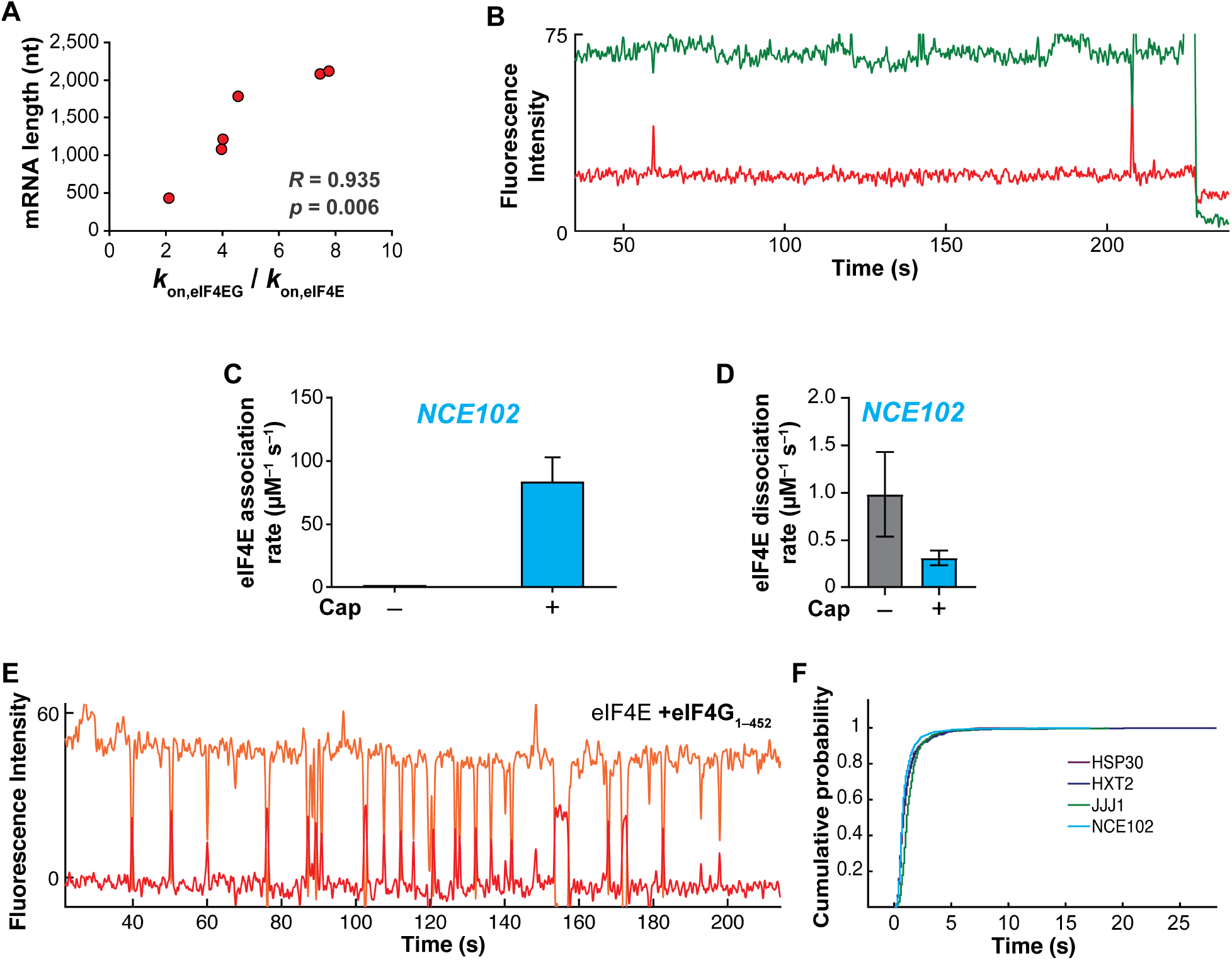
Relates to Figure 2. eIF4E–mRNA kinetics with eIF4G1, eIF4G1_1–452_ and uncapped mRNA. **A.** Correlation of eIF4G1 stimulation of eIF4E-mRNA association with mRNA length. **B.** Representative smFRET trace from an experiment with uncapped *NCE102* and eIF4E•eIF4G1. **C.** eIF4E-mRNA association rate on uncapped *vs.* capped *NCE102*. **D.** eIF4E-mRNA dissociation rate on uncapped *vs.* capped *NCE102*. **E.** Sample smFRET trajectory for eIF4E–mRNA binding in the presence of eIF4G1_1–452_. **F.** Cumulative distribution function of the eIF4E-mRNA interaction lifetimes in the presence of eIF4G1_1–452_.

**Supplemental Figure 4.**
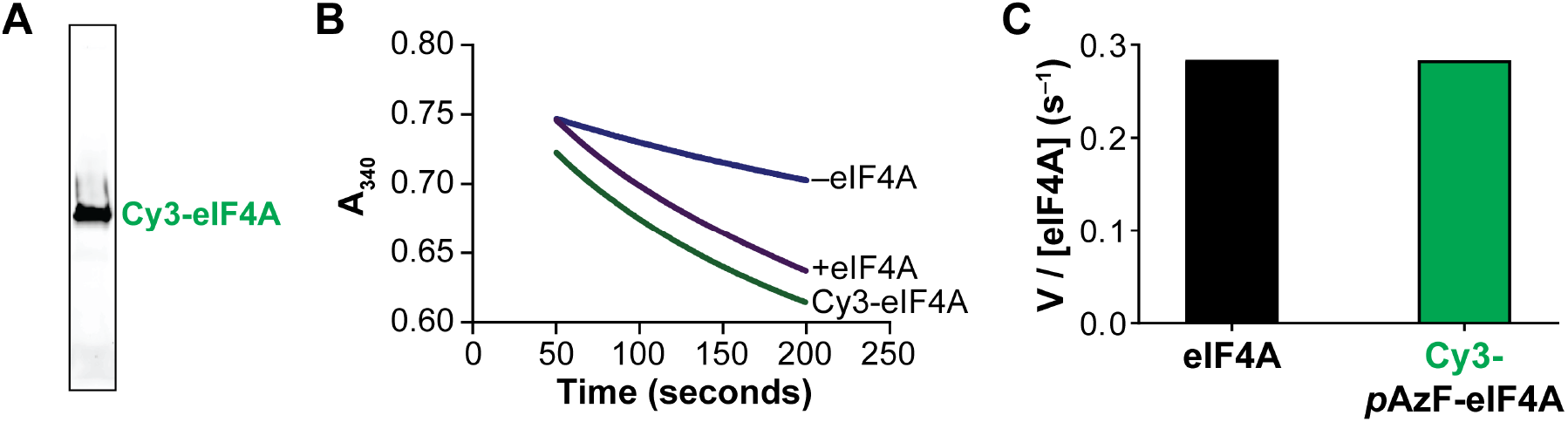
Relates to Figure 5. Preparation and validation of Cy3-eIF4A. **A.** SDS-PAGE analysis of Cy3-eIF4A, imaged for Cy3 fluorescence. **B.** Time-courses of ATP hydrolysis catalyzed by eIF4A and Cy3-eIF4A in the presence of poly(U) RNA and eIF4G1, monitored at 340 nm in an NADH-coupled assay, and compared with a no-enzyme control reaction. **C.** Quantitation of specific activity for eIF4A and Cy3-eIF4A from the time-course data in panel B.

**Supplemental Figure 5.**
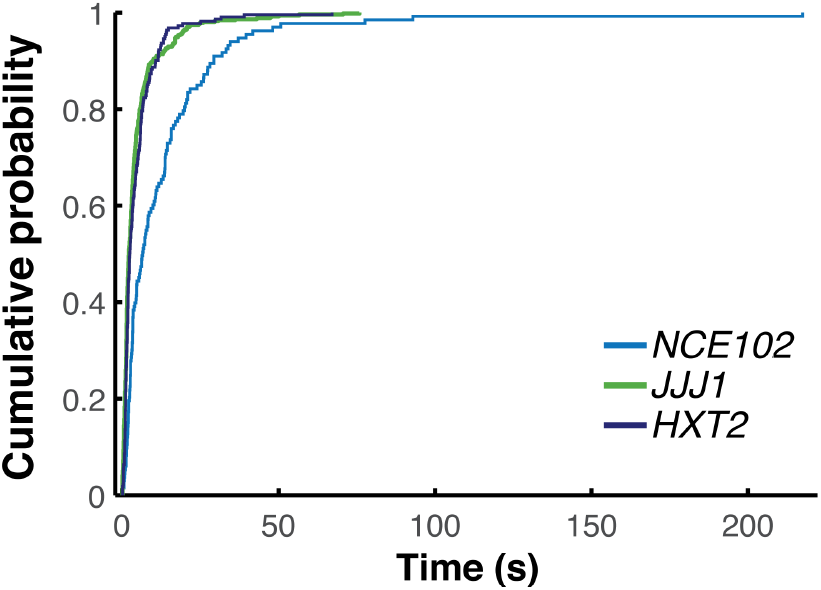
Relates to Figure 5. Kinetics of free eIF4A•mRNA dissociation. **A.** Cumulative distribution functions for lifetimes of transient free eIF4A–mRNA binding events, indicating double-exponential behavior.

## STAR METHODS

### RESOURCE AVAILABILITY

#### Lead contact

The lead contact for this study is Seán E. O’Leary.

#### Materials Availability

All materials and reagents described in this study are available on request without restriction. Further information and requests for resources and reagents should be directed to and will be fulfilled by the lead contact.

#### Data and code availability

The published article includes all relevant data generated or analyzed during the study. Source data for all figures are available in the Supplemental Information. This study did not generate any code.

### EXPERIMENTAL MODEL AND SUBJECT DETAILS

Genes encoding translation factors were expressed from pET-28a(+) (Qiagen) or pTYB2 (New England Biolabs). Overexpression was carried out at 37 °C in LB medium, in volumes ranging from 1 L to 12 L. *E. coli* BL21(DE3) CodonPlus RIL or BL21(DE3) cells expressing the target recombinant protein were grown to an OD_600_ of 0.5 – 1 at 37 °C. Overexpression was then induced *via* addition of 0.5 mM IPTG, then the overexpression was allowed to proceed overnight at 16 °C. For eIF4G, the induction was carried out at 37 °C for 2 – 3 hours. The resulting cells were harvested and stored at –80 °C until purification.

### METHOD DETAILS

#### In vitro transcription, RNA processing and labeling of oligonucleotides

DNA templates to be transcribed were PCR-amplified from Yeast Genomic DNA (EMD Millipore) with Phusion DNA polymerase (NEB) using standard procedures. Primers sequences are reported in Supplemental Table 11. A T7 promoter (TAATACGACTCACTATA<u>GG)</u> was incorporated into the PCR product through the forward primer, where the underlined bases become +1 and +2 nucleotides added to the transcript. The resulting templates were *in-vitro* transcribed to produce RNA using in-house purified or commercial (NEB) T7 RNA. Transcription reactions were typically carried out on a 40 µL scale, with ∼5 µg of DNA template, in a buffer consisting of 200 mM Tris-HCl, pH 7.9, 0.05% (v/v) Triton-X-100, 15 mM spermidine, 2 mM each NTP and 5 mM DTT. MgCl_2_ concentrations in this buffer were optimized for each transcript by titration, and were 15 – 30 mM. Unreacted nucleotides were removed from the transcription mixture with MicroBio-Spin gel filtration columns (P30, Bio-Rad), and the RNA product was then precipitated using ½ volumes of 7.5M lithium chloride. The resulting RNA pellet was redissolved after three washes with 80% EtOH and run on a native 1% TBE-Agarose gel, to check for integrity. The RNAs were then capped using the ScriptCap™ m^7^G Capping System (CellScript) with the following modifications to the manufacturer’s protocol: the incubation time with the capping enzyme was increased to two hours, and the volume of enzyme added was increased twofold. A poly(A) tail was added to the mRNA immediately after capping, using *E. coli* poly(A) polymerase (NEB), following the manufacturer’s guidelines. The capped and tailed mRNA was then re-purified by organic extraction with acidic phenol-chloroform, precipitated with 0.1 volumes of 3 M sodium acetate and two volumes of ethanol, and resuspended in RNase-free water.

5ʹ biotinylated and 3ʹ amino-modified oligonucleotides purchased from IDT were reacted with a 1:8 molar ratio of oligonucleotide to NHS-ester derivatives of Cy5 (for intramolecular FRET experiments with dual labeled RNA), Cy3 and Cy3.5 in 0.1 M sodium bicarbonate for four hours at room temperature, followed by four successive chloroform extractions to remove unreacted dye, and by buffer-exchange into ddH_2_O using MicroBio-Spin gel filtration columns (P30, Bio-Rad). Labeling efficiency was typically 50 – 75% as measured by UV/visible spectrophotometry. The labeled oligonucleotides were stored at –20 °C and used without further purification.

#### Protein purification and lab

His_6_-tagged yeast eIF4E(A124C) was purified as described previously (O’Leary et al., 2013). Briefly, cell lysate from the overexpression culture was loaded onto a gravity-flow Ni-NTA agarose column (Qiagen) pre-equilibrated with eIF4E buffer (50 mM HEPES-KOH, pH 7.4, 150 mM KCl, 2.5 mM TCEP). The column was then washed with 40 column volumes of eIF4E buffer containing 40 mM imidazole to remove nonspecifically-bound proteins, then eIF4E was eluted using eIF4E buffer containing 250 mM imidazole. Imidazole was removed from the protein eluate by desalting on a Bio-Rad 10-DG column equilibrated in containing 50 mM HEPES-KOH, pH 7.4, 150 mM KCl, 0.5 mM TCEP. The resulting protein was immediately labeled with a sulfonated Cyanine 5 maleimide (Lumiprobe) overnight at 6 °C in darkness. Unreacted fluorophore was then removed by desalting on a BioRad 10-DG column. The labeled protein was purified by gel filtration using a Superdex 75 Increase column (GE healthcare), equilibrated in storage buffer (50 mM HEPES-KOH, pH 7.4, 150 mM KCl, 2.5 mM TCEP). The labeling efficiency was assessed by UV/visible spectrophotometry, and was typically ∼50%. The protein was stored at 6 °C in darkness, and was prepared freshly every week as needed.

The plasmid containing recombinant full-length eIF4G1 was a gift from Sarah Walker (also available on Addgene as plasmid #122248). This full-length eIF4G1 construct with a C-terminal chitin binding domain fusion was purified according to a published procedure (Liu et al., 2019), with the following modifications: the cells were lysed using a sonicator, and the final protein after elution was stored in 250 mM KCl instead of 250 mM KOAc, skipping the dialysis after the anion-exchange chromatography step in the procedure, and DTT for storage was substituted with 2.5 mM TCEP. Briefly, *E. coli* cells expressing full-length eIF4G1 were thawed, and lysed using a sonicator after resuspending in Intein Lysis Buffer (50 mM HEPES-KOH, pH 7.4, 500 mM KCl, 1 mM EDTA). The lysate was then clarified by centrifugation at 20,000 × *g* for 15 minutes. The clarified lysate was rocked with 4 mL of chitin resin (New England Biolabs) for 30 minutes at 4 °C. The resin was then loaded into a gravity-flow column and washed with 100 mL of Intein Lysis Buffer. The column was then treated with micrococcal nuclease to remove nucleic acids from *E. coli* which co-purified with eIF4G. Briefly, the resin was first equilibrated with microccocal nuclease buffer (50 mM HEPES-KOH, pH 7.4, 100 mM KCl, 2 mM CaCl_2_). Then, a 3 mL solution containing 3 U/µL microccocal nuclease was passed through the column. The resin was then incubated for 30 minutes at 37 °C. Following nuclease treatment, the column was washed with a further 50 mL of lysis buffer. The column was then flushed with 8 mL of lysis buffer containing 50 mM DTT, and 6 mL was allowed to pass through the column, which was then sealed and incubated overnight at 6 °C. The following day, the cleaved protein was eluted with 10 mL of lysis buffer. The resulting protein solution was diluted to 100 mM KCl with lysis buffer lacking KCl, then manually loaded onto a Q HP column (1 mL; GE Healthcare Life Sciences) equilibrated in 50 mM HEPES-KOH, pH 7.4, 10% (v/v) glycerol, 2.5 mM TCEP, 100 mM KCl, and washed with five column volumes of buffer containing 50 mM HEPES-KOH, pH 7.4, 10% (v/v) glycerol, 2.5 mM TCEP, 100 mM KCl. The column was then eluted manually with a step-gradient of 150, 200, 250 mM KCl in 50 mM HEPES-KOH, pH 7.4, 10% (v/v) glycerol, 2.5 mM TCEP (one column volume for each step). Eluate fractions were analyzed by SDS-PAGE; eIF4G typically eluted above ∼220 mM KCl. Single-use aliquots of purified eIF4G1 were prepared and were stored at –80 °C. The His_6_-tagged eIF4G_1–452_ fragment (an *NdeI* fragment of the eIF4G1 CDS) was purified essentially as for eIF4E, but 1 M KCl was included in the purification buffers.

His_6_-tagged recombinant eIF4A was purified as described previously (O’Leary et al., 2013). *E. coli* cells expressing recombinant eIF4A were first thawed and resuspended in eIF4A lysis buffer (50 mM HEPES-KOH, pH 7.4, 300 mM KCl, 2.5 mM TCEP). After sonication for cell lysis, the resulting lysate was clarified by spinning at 20,000 × *g* for 15 minutes. The clarified lysate was applied to Ni-NTA agarose (equilibrated in lysis buffer) as a first step, after filtering the lysate through a 0.22 µm syringe filter. The bound protein was eluted with lysis buffer containing 250 mM imidazole after washing with 10 column volumes of lysis buffer containing 40 mM Imidazole. The eluate was buffer-exchanged to Buffer A using a BioRad 10-DG column (50 mM HEPES-KOH, pH 7.4, 100 mM KCl, 2.5 mM TCEP) and subjected to anion-exchange chromatography using a 5 mL Q HP anion-exchange column (GE Healthcare). The column was eluted with a linear gradient of 0.1 – 1 M KCl. eIF4A typically eluted at 250 mM KCl. Fractions containing eIF4A were identified by SDS-PAGE analysis, then pooled, concentrated by centrifugal ultrafiltration, and further purified by gel filtration chromatography using a Superdex 200 column (GE healthcare) equilibrated in 50 mM HEPES-KOH, pH 7.4, 100 mM KOAc, 2.5 mM TCEP, 10% (v/v) glycerol. The final protein sample was divided into single-use aliquots and stored in storage buffer at –80 °C.

For preparation of labeled eIF4A, a construct was designed that expresses the native eIF4A sequence with an N-terminal Met-Ala-(*p*Az)Phe tripeptide extension for unnatural amino acid incorporation. This plasmid was co-transformed into *E. coli* BL21(DE3) cells with the pEVOL-ps plasmid (a generous gift from Abhishek Chatterjee, Boston College) under dual selection with chloramphenicol and kanamycin. The resulting transformants were grown in a 10 mL starter culture overnight. Afterwards, a 1 L LB medium flask was inoculated with the starter culture and grown to an O.D._600_ value of 0.5. 1 mM 4-azidophenylalanine was then added to the culture medium along with 2 mM arabinose to induce tRNA/aminoacyl-tRNA synthetase expression. Finally, 2 mM IPTG was added to induce expression of eIF4A. Overexpression was allowed to proceed for 5 hours in darkness, to avoid photochemical damage to the unnatural amino acid. The cells were harvested and stored at –80 °C until purification. MA(*p*AzF)-eIF4A was purified identically to unlabeled recombinant eIF4A, with the exception that after initial Ni-NTA purification the protein was treated with DBCO-Cy3 overnight to conjugate the fluorophore to the unnatural amino acid.

#### Steady-state ATPase assay for eIF4A activity

The NADH-coupled ATPase assay was carried out according to a published procedure (Bradley and De La Cruz, 2012). Briefly, reactions were assembled on ice and started by adding Mg-ATP. The reaction was set up with the KMg75 buffer (20 mM HEPES-KOH, pH 7.4, 75 mM KCl, 1 mM DTT, 5 mM MgCl_2_), 250 nM eIF4A or Cy3-eIF4A, 125 nM full-length eIF4G, 1 mM ATP, 1 mM (measured as concentration of bases) poly(U) RNA (Sigma), lactate dehydrogenase (20 U/mL final concentration), and pyruvate kinase (100 U/mL) Absorbance was recorded at 340 nM with a Shimadzu UV2600 UV-visible spectrophotometer, measuring the decrease of NADH absorbance with time. The slope of the absorbance *vs*. time graph was converted to the rate of ATP hydrolysis using an extinction coefficient of 6,220 M^−1^ cm^−1^ for NADH, and normalized to the eIF4A concentration to yield *V* / *E*_0_. Control reactions to establish the background rate of NADH oxidation included no eIF4A and no RNA.

#### Single-molecule experiments

The custom RS instrument was set up as described previously (Chen et al., 2014; Çetin *et al*., 2020). The RNA to be immobilized was hybridized through its poly(A) tail to (dT)_45_ conjugated to biotin at its 5ʹ end and Cy3 or Cy3.5 at its 3ʹ end, to act as a FRET donor. Annealing was performed using a thermocycler, by heating 100 nM labeled oligonucleotide to 98 °C for two minutes in the presence of two- to five-fold molar excess of mRNA, followed by cooling to 4 °C at a ramp speed of 0.1 °C s^−1^. The resulting mRNA:(dT)_45_ duplex was diluted to 3-10 nM fluorophore in smFRET assay buffer prior to immobilization on the ZMW using the assay buffer.

Zero-mode waveguides were set up as described previously (Chen et al. 2014). Briefly, the ZMW chip was hydrated with assay buffer (final concentrations of 50 mM HEPES-KOH, pH 7.4, 3 mM Mg(OAc)_2_, 100 mM KOAc) for two minutes, followed by incubation with 16 µM NeutrAvidin (Thermo Scientific) for five minutes to allow immobilization of biomolecules. The chip was then washed three times with the assay buffer, followed by addition of 10 nM mRNA:biotin-(dT)_45_-Cy3.5 duplex, which was allowed to immobilize for 20 minutes. The chip was then washed again three times to remove non-immobilized nucleic acids, and an imaging buffer containing PCA/PCD oxygen scavenging system and photostabilizer (TSY) (Aitken et al., 2008, Chen et al. 2014) was added. Prior to imaging on the RS II, the chip was treated with 5% (v/v) each of BioLipidure 203 and 206, 1 mg/mL BSA and unlabeled eIF4E; this blocking step mitigates non-specific Cy5-eIF4E interactions with the surface. Inclusion of this step did not detectably alter the kinetics of eIF4E-mRNA interaction. After initiating the imaging on the RS II, between 4 and 30 nM Cy5-eIF4E were robotically injected onto the waveguide, starting the binding reaction. The ZMWs were imaged with 10-minute movies acquired at 10 frames/second, at 0.7 µW/µm^2^ green (532 nm) laser power and 0.07 µW/µm^2^ red (642 nm) laser power (for dual illumination experiments).

#### Single-molecule data processing and analysis

Raw movie data were extracted and analyzed with an in-house MATLAB processing pipeline as described previously (Chen et al. 2014; O’Leary *et al*., 2013; Çetin *et al*., 2020). Image files were first converted to fluorescence *vs.* time traces. The locations of events in the traces were then manually assigned, resulting in distributions of event and inter-event durations (i.e., lifetimes and arrival times).

For the two-color FRET experiments, events showing anticorrelated bursts between the FRET donor and acceptor were manually assigned as FRET. For the three-color FRET experiments, events showing Cy3 signal only were scored as free eIF4A binding. Events showing appearance of Cy3 fluorescence with a concomitant increase in Cy5 fluorescence significantly beyond the negligible expected bleedthrough (Chen *et al*., 2014), and showing apparent FRET efficiency changes during the ensuing binding event, were characterized as eIF4E-eIF4A FRET. A further type of binding event where eIF4E-eIF4A FRET disappeared while eIF4A stayed bound was also scored in both the number of occurrences and the dwell time of the initial FRET event. During these Cy3 fluorescence pulses, the dwell time of FRET when eIF4E-eIF4A FRET reappeared was also quantified. Rate constants were quantified by exponential fitting as described above.

### QUANTIFICATION AND STATISTICAL ANALYSIS

Numbers of independent replicates, numbers of molecules analyzed in each replicate experiment, and goodness-of-fit metrics for reported rate constants (Figures 1 – 6; Supplemental Figure 3) are reported in Supplemental Tables 2 – 10. For Figures 5B and 6B, *n* is the number of single molecules analyzed to enumerate distinct event types.

For kinetic analysis, arrival-time or lifetime distributions for two-color smFRET experiments were constructed from analysis of events occurring on at least 100 mRNA molecules, which included at least 500 events, and typically more than 1,000 events. Addition to the analysis of further molecules beyond this number neither significantly altered the kinetic parameters obtained, nor improved the quality of data fitting. Empirical cumulative distribution functions for unbinned distributions were fit in MATLAB, using nonlinear least-squares regression, to either single-exponential *P*(*t*) = 1 − *Ae*^−*kt*^ or double-exponential *P*(*t*) = *A* (1 − *e*^−*k*_1_*t*^) + (1 − *A*)(1 − *e*^−*k*_2_*t*^) models, as appropriate. For double-exponential arrival-time distributions, the fast-phase rate, which typically constituted > 70% of the amplitude, was used for comparison of eIF4E binding between different mRNAs and conditions.

For goodness-of-fit evaluation, fits typically had an *R*^2^ value > 0.99 (for all distributions generated from experiments with eIF4E-G, eIF4F) and greater than 0.95 (for experiments containing only eIF4E and eIF4E-eIF4A.). Root-mean-squared errors of the fits for arrival-time distributions were typically 0.02 or a lower value; lifetime distributions showed more variable RMSE values with an upper limit of 0.1. RMSE values and 95% confidence intervals of the fits are provided in the Supplemental Tables.

For correlation of kinetic parameters with mRNA lengths (Figure 1G; Supplemental Figure 1D,E; Figure 2B) Pearson correlation coefficients (*R*) were calculated using GraphPad Prism software (Version 9.1.). Correlations with *p* < 0.05 based on a two-tailed Student’s *t*-test were considered significant.

## Notes

### Competing Interest Statement

The authors have declared no competing interest.

