## Supplemental Information for "mRNA- and factor-driven dynamic variability controls eIF4F-cap recognition for translation initiation"

**TABLE OF CONTENTS**


**Supplemental Tables** 3

**Supplemental Table 1. Relates to Figure 1.** 3
mRNA constructs

**Supplemental Table 2. Relates to Figure 1** 4
Single-molecule kinetic data for eIF4E–mRNA interactions

**Supplemental Table 3. Relates to Figure 2.** 5
Single-molecule kinetic data for eIF4E–mRNA interactions in the presence of eIF4G1

**Supplemental Table 4. Relates to Figure 3.** 6
Single-molecule kinetic data for eIF4E–mRNA interactions in the presence of free eIF4A and ATP

**Supplemental Table 5. Relates to Figure 3.** 7
Single-molecule kinetic data for eIF4E–mRNA interactions in the presence of free eIF4A and ATP-$\gamma$-S

**Supplemental Table 6. Relates to Figure 4.** 8
Single-molecule kinetic data for eIF4E–mRNA interactions in the presence of eIF4F without ATP

**Supplemental Table 7. Relates to Figure 4.** 9
Single-molecule kinetic data for eIF4E–mRNA interactions in the presence of eIF4F and ATP

**Supplemental Table 8. Relates to Figure 5.** 10
eIF4A•mRNA association rates for all binding-event types, and dissociation rates for mRNA complexes with free eIF4A

**Supplemental Table 9. Relates to Figure 5.** 11
Dissociation rates for eIF4A–mRNA binding events showing eIF4A–eIF4E FRET

**Supplemental Table 10. Relates to Figure 6.** 12
eIF4E–mRNA dissociation rates following initial eIF4F–mRNA binding event and eIF4E mRNA rebinding.

**Supplemental Table 11. Relates to STAR Methods and Key Resources.** 13
Primers for mRNA template amplification from yeast genomic DNA

**Supplemental Discussion** 14

**Supplemental Discussion D1** 14
Accuracy of association rate constants from smFRET experiment

**Supplemental Table 1**. **Relates to Figure 1.** mRNA constructs.

| Gene | Length (nt) | | | | 5' UTR Annotation* | 3' UTR Annotation* |
|---|---|---|---|---|---|---|
| | 5' UTR | CDS | 3' UTR | Total | | |
| *JJJ1* | 49 | 1773 | 263 | 2085 | Yassour | Yassour |
| *HSP30* | 77 | 999 | 140 | 1216 | Nagalakshmi | Nagalakshmi |
| *NCE102* | 171 | 522 | 387 | 1080 | Nagalakshmi | Nagalakshmi |
| *HXT2* | 40 | 1626 | 139 | 1805 | Nagalakshmi | Nagalakshmi |
| *MIM1* | 24 | 342 | 68 | 434 | Nagalakshmi | Nagalakshmi |
| *POP5* | 7 | 522 | 74 | 605 | Yassour | Nagalakshmi |

* Annotations from: Yassour *et al.* (2009); Nagalakshmi *et al.* (2008)

**Supplemental Table 2. Relates to Figure 1.** Single-molecule kinetic data for eIF4E–mRNA interactions.

| mRNA | Repl. (No. of Mol.) | $k_{on}$ ($\mu M^{-1}$ $s^{-1}$) | | | | $k_{off}$ ($s^{-1}$) | | | |
|---|---|---|---|---|---|---|---|---|---|
| | | Fitted rate | 95% CI | | RMSE | Fitted Rate | 95% CI | | RMSE |
| | | | Low | High | | | Low | High | |
| *JJJ1* | 1 (100) | 7.4 | 7.07 | 7.68 | 0.02 | 0.33 | 0.31 | 0.35 | 0.08 |
| | 2 (100) | 7.5 | 7.25 | 7.69 | 0.02 | 0.56 | 0.53 | 0.61 | 0.07 |
| | 3 (100) | 4.2 | 4.14 | 4.26 | 0.02 | 0.41 | 0.38 | 0.43 | 0.06 |
| *NCE102* | 1 (322) | 21.9 | 19.5 | 21.0 | 0.01 | 0.418 | 0.38 | 0.45 | 0.05 |
| | 2 (100) | 25.0 | 23 | 27.3 | 0.02 | 0.53 | 0.56 | 0.59 | 0.08 |
| | 3 (100) | 20.6 | 22.1 | 22.8 | 0.02 | 0.358 | 0.35 | 0.37 | 0.03 |
| *HXT2* | 1 (100) | 4.9 | 4.59 | 5.35 | 0.02 | 1.06 | 0.91 | 1.2 | 0.10 |
| | 2 (100) | 8.3 | 7.42 | 9.72 | 0.02 | 1.02 | 0.93 | 1.1 | 0.05 |
| | 3 (100) | 14.9 | 13.9 | 15.1 | 0.02 | 0.56 | 0.52 | 0.59 | 0.07 |
| *HSP30* | 1 (100) | 18.4 | 17.9 | 18.9 | 0.02 | 0.91 | 0.85 | 0.97 | 0.09 |
| | 2 (100) | 27.5 | 24.5 | 30.4 | 0.02 | 1.0 | 0.91 | 1.17 | 0.05 |
| | 3 (100) | 24.0 | 21.3 | 26.8 | 0.01 | 0.73 | 0.67 | 0.77 | 0.05 |

**Supplemental Table 3. Relates to Figure 2.** Single-molecule kinetic data for eIF4E–mRNA interactions in the presence of eIF4G1.

| mRNA | Repl. (No. of mol.) | $k_{on}$ ($\mu M^{-1}$ $s^{-1}$) | | | | $k_{off,fast}$ ($s^{-1}$) | | | $k_{off,slow}$ ($s^{-1}$) | | | Slow phase ampli-tude (%) |
|---|---|---|---|---|---|---|---|---|---|---|---|---|
| | | Fitted Rate | 95% CIs | | RMSE | Fitted Rate | 95% CIs | | Fitted Rate | 95% CIs | | |
| | | | Low | High | | | Low | High | | Low | High | |
| *JJJ1* | 1 (118) | 50.1 | 48.0 | 51.9 | 0.01 | 0.25 | 0.24 | 0.26 | 0.038 | 0.033 | 0.043 | 17 |
| | 2 (106) | 41.8 | 40.1 | 43.6 | 0.02 | 0.55 | 0.53 | 0.57 | 0.079 | 0.058 | 0.099 | 21 |
| *NCE102* | 1 (126) | 107.1 | 104.5 | 109.7 | 0.02 | 0.39 | 0.39 | 0.40 | 0.046 | 0.043 | 0.049 | 23 |
| | 2 (100) | 70.6 | 68.9 | 72.3 | 0.01 | 0.32 | 0.31 | 0.33 | 0.032 | 0.013 | 0.051 | 17 |
| *HXT2* | 1 (100) | 48.6 | 48.6 | 50.5 | 0.02 | 0.36 | 0.35 | 0.37 | 0.046 | 0.042 | 0.049 | 16 |
| | 2 (123) | 34.2 | 33.1 | 35.3 | 0.02 | 0.34 | 0.33 | 0.35 | 0.023 | 0.020 | 0.026 | 16 |
| *HSP30* | 1 (105) | 83.3 | 81.3 | 85.3 | 0.01 | 0.77 | 0.74 | 0.80 | 0.11 | 0.10 | 0.12 | 20 |
| | 2 (100) | 75.4 | 72.8 | 77.9 | 0.006 | 0.63 | 0.61 | 0.64 | 0.090 | 0.087 | 0.094 | 27 |

**Supplemental Table 4. Relates to Figure 3.** Single-molecule kinetic data for eIF4E–mRNA interactions in the presence of free eIF4A and ATP.

| mRNA | Repl. (No. of Mol.) | $k_{on}$ ($\mu M^{-1}$ $s^{-1}$) | | | | $k_{off}$ ($s^{-1}$) | | | |
|---|---|---|---|---|---|---|---|---|---|
| | | Fitted rate | 95% CI | | RMSE | Fitted Rate | 95% CI | | RMSE |
| | | | Low | High | | | Low | High | |
| *JJJ1* | 1 (117) | 29 | 27.9 | 29.1 | 0.01 | 1.33 | 1.17 | 1.49 | 0.07 |
| | 2 (103) | 24.8 | 24.1 | 25.5 | 0.01 | 1.44 | 1.23 | 1.63 | 0.03 |
| *NCE102* | 1 (114) | 59.9 | 58.2 | 61.8 | 0.01 | 0.81 | 0.77 | 0.85 | 0.03 |
| | 2 (100) | 80.4 | 71.2 | 89.9 | 0.02 | 0.86 | 0.78 | 0.95 | 0.04 |
| *HXT2* | 1 (110) | 24.7 | 23.6 | 25.8 | 0.01 | 0.69 | 0.65 | 0.73 | 0.04 |
| | 2 (106) | 24.9 | 23.8 | 25.9 | 0.01 | 0.76 | 0.72 | 0.80 | 0.04 |
| *HSP30* | 1 (103) | 30 | 29.4 | 30.6 | 0.01 | 1.25 | 1.13 | 1.36 | 0.04 |
| | 2 (100) | 25.7 | 23.4 | 27.0 | 0.01 | 0.89 | 0.83 | 0.95 | 0.04 |

**Supplemental Table 5. Relates to Figure 3.** Single-molecule kinetic data for eIF4E–mRNA interactions in the presence of free eIF4A and ATP-γ-S.

| mRNA | Repl. (No. of Mol.) | $k_{on}$ ($\mu M^{-1}$ $s^{-1}$) | | | | $k_{off}$ ($s^{-1}$) | | | |
|---|---|---|---|---|---|---|---|---|---|
| | | Fitted rate | 95% CI | | RMSE | Fitted Rate | 95% CI | | RMSE |
| | | | Low | High | | | Low | High | |
| *JJJ1* | 1 (100) | 29.4 | 27.1 | 31.7 | 0.03 | 1.4 | 0.1 | 1.7 | 0.11 |
| | 2 (100) | 19.3 | 18.4 | 23.2 | 0.01 | 1.2 | 1.1 | 1.3 | 0.05 |
| *NCE102* | 1 (108) | 52.6 | 50.6 | 54.5 | 0.01 | 1.62 | 1.4 | 1.84 | 0.08 |
| | 2 (100) | 36.9 | 33. | 40.2 | 0.02 | 0.98 | 0.89 | 1.07 | 0.07 |

**Supplemental Table 6. Relates to Figure 4.** Single-molecule kinetic data for eIF4E–mRNA interactions in the presence of eIF4F without ATP.

| mRNA | Repl. (No. of mol.) | $k_{on}$ ($\mu M^{-1}$ $s^{-1}$) | | | | $k_{off,fast}$ ($s^{-1}$) | | | $k_{off,slow}$ ($s^{-1}$) | | | | Slow phase amplitude (%) |
|---|---|---|---|---|---|---|---|---|---|---|---|---|---|
| | | Fitted Rate | 95% CIs | | RMSE | Fitted Rate | 95% CIs | | Fitted Rate | 95% CIs | | RMSE | |
| | | | Low | High | | | Low | High | | Low | High | | |
| | 1 (104) | 33.5 | 32.7 | 34.3 | 0.01 | 0.27 | 0.26 | 0.28 | 0.068 | 0.054 | 0.073 | 0.01 | 32 |
| *JJJ1* | 2 (119) | 53.1 | 51.21 | 55.0 | 0.02 | 0.28 | 0.27 | 0.28 | 0.043 | 0.038 | 0.047 | 0.01 | 18 |
| | 1 118 | 75.8 | 72.8 | 78.7 | 0.02 | 0.45 | 0.43 | 0.46 | 0.076 | 0.067 | 0.086 | 0.01 | 20 |
| *NCE102* | 2 (121) | 60.0 | 58.58 | 61.6 | 0.01 | 0.37 | 0.36 | 0.38 | 0.077 | 0.072 | 0.082 | 0.01 | 27 |
| | 1 110 | 53.5 | 55.4 | 51.4 | 0.02 | 0.45 | 0.43 | 0.46 | 0.053 | 0.049 | 0.057 | 0.02 | 21 |
| *HXT2* | 2 (98) | 45.6 | 42.5 | 48.7 | 0.02 | 0.36 | 0.34 | 0.38 | 0.054 | 0.027 | 0.081 | 0.02 | 29 |
| | 1 (110) | 101.7 | 97.7 | 105.7 | 0.01 | 0.64 | 0.61 | 0.68 | 0.097 | 0.089 | 0.11 | 0.02 | 30 |
| *HSP30* | 2 (160) | 102.7 | 99 | 106.4 | 0.01 | 0.63 | 0.62 | 0.65 | 0.083 | 0.081 | 0.086 | 0.02 | 34 |

**Supplemental Table 7. Relates to Figure 4.** Single-molecule kinetic data for eIF4E–mRNA interactions in the presence of eIF4F and ATP.

| mRNA | Repl. (No. of mol.) | $k_{on}$ ($\mu M^{-1}$ $s^{-1}$) Fitted Rate | 95% CIs Low | 95% CIs High | RMSE | $k_{off,fast}$ ($s^{-1}$) Fitted Rate | 95% CIs Low | 95% CIs High | $k_{off,slow}$ ($s^{-1}$) Fitted Rate | 95% CIs Low | 95% CIs High | RMSE | Slow phase amplitude (%) |
|---|---|---|---|---|---|---|---|---|---|---|---|---|---|
| *JJ1* | 1 (192) | 43.3 | 42.3 | 44.3 | 0.01 | 0.71 | 0.68 | 0.73 | 0.099 | 0.096 | 0.10 | 0.01 | 30 |
| | 2 (280) | 32.3 | 31.6 | 33.1 | 0.01 | 0.69 | 0.67 | 0.71 | 0.098 | 0.095 | 0.10 | 0.01 | 24 |
| *NCE102* | 1 (94) | 83.6 | 81.7 | 85.6 | 0.01 | 0.99 | 0.94 | 1.0 | 0.088 | 0.076 | 0.10 | 0.01 | 29 |
| | 2 (118) | 71.7 | 70.2 | 73.2 | 0.01 | 0.74 | 0.71 | 0.76 | 0.083 | 0.078 | 0.089 | 0.01 | 30 |
| *HXT2* | 1 (307) | 38.5 | 37.5 | 39.4 | 0.02 | 0.74 | 0.71 | 0.76 | 0.117 | 0.11 | 0.13 | 0.02 | 17 |
| | 2 (116) | 36.83 | 35.9 | 37.7 | 0.02 | 0.66 | 0.63 | 0.68 | 0.098 | 0.092 | 0.10 | 0.02 | 32 |
| *HSP30* | 1 (100) | 26.1 | 25.2 | 27.03 | 0.01 | 0.85 | 0.75 | 0.94 | 0.19 | 0.11 | 0.28 | 0.03 | 16 |
| | 2 (151) | 39.5 | 37.2 | 41.8 | 0.01 | 0.99 | 0.91 | 1.1 | 0.14 | 0.079 | 0.19 | 0.02 | 16 |

**Supplemental Table 8. Relates to Figure 5.** eIF4A•mRNA association rates for all binding-event types, and dissociation rates for mRNA complexes with free eIF4A.

| mRNA | Repl. (No. of mol.) | $k_{on}$ ($\mu M^{-1}$ $s^{-1}$) | | | | $k_{off,fast}$ ($s^{-1}$) | | | $k_{off,slow}$ ($s^{-1}$) | | | | Slow phase amplitude (%) |
|---|---|---|---|---|---|---|---|---|---|---|---|---|---|
| | | Fitted Rate | 95% CIs | | RMSE | Fitted Rate | 95% CIs | | Fitted Rate | 95% CIs | | RMSE | |
| | | | Low | High | | | Low | High | | Low | High | | |
| *JJ1* | 1 (85) | 2.41 | 2.3 | 2.5 | 0.01 | 0.37 | 0.35 | 0.38 | 0.05 | 0.04 | 0.07 | 0.01 | 10 |
| | 2 (100) | 3.1 | 3.0 | 3.3 | 0.01 | 0.15 | 0.14 | 0.15 | 0.02 | 0.01 | 0.03 | 0.01 | 12 |
| *NCE102* | 1 (92) | 3.6 | 3.1 | 4.1 | 0.03 | 0. 21 | 0.15 | 0.3 | 0.06 | 0.05 | 0.07 | 0.03 | 61 |
| | 2 (100) | 4 | 3.6 | 4.5 | 0.02 | 0.70 | 0.55 | 0.85 | 0.13 | 0.12 | 0.14 | 0.02 | 75 |
| *HXT2* | 1 (100) | 2.5 | 2.3 | 2.7 | 0.02 | 0.3 | 0.24 | 0.36 | 0.09 | 0 | 0.20 | 0.02 | 15 |
| | 2 (102) | 5.6 | 4.7 | 6.4 | 0.02 | 0.35 | 0.28 | 0.41 | 0.088 | 0.02 | 0.15 | 0.10 | 45 |

**Supplemental Table 9. Relates to Figure 5.** Dissociation rates for eIF4A–mRNA binding events showing eIF4A–eIF4E FRET.

| mRNA | Repl. (No. of Mol.) | $k_{off}$ ($s^{-1}$) Fitted Rate | 95% CI Low | 95% CI High | RMSE |
|---|---|---|---|---|---|
| *JJJ1* | 1 (85) | 0.033 | 0.033 | 0.034 | 0.05 |
| | 2 (100) | 0.02 | 0.02 | 0.02 | 0.02 |
| *NCE102* | 1 (92) | 0.036 | 0.035 | 0.038 | 0.05 |
| | 2 (100) | 0.045 | 0.04 | 0.05 | 0.06 |
| *HXT2* | 1 (100) | 0.037 | 0.035 | 0.039 | 0.03 |
| | 2 (102) | 0.028 | 0.027 | 0.031 | 0.05 |

**Supplemental Table 10. Relates to Figure 6.** eIF4E–mRNA dissociation rates following initial eIF4F–mRNA binding event and eIF4E–mRNA rebinding.

| mRNA | Repl. (No. of Mol.) | $k_{off,\ initial}$ ($s^{-1}$) | | | | $k_{off,\ rebinding}$ ($s^{-1}$) | | | |
|---|---|---|---|---|---|---|---|---|---|
| | | Fitted rate | 95% CI | | RMSE | Fitted Rate | 95% CI | | RMSE |
| | | | Low | High | | | Low | High | |
| *JJJ1* | 1 (85) | 0.05 | 0.05 | 0.06 | 0.04 | 0.099 | 0.093 | 0.10 | 0.04 |
| | 2 (100) | 0.08 | 0.08 | 0.09 | 0.06 | 0.12 | 0.11 | 0.12 | 0.06 |
| *NCE102* | 1 (92) | 0.10 | 0.09 | 0.11 | 0.06 | 0.10 | 0.09 | 0.10 | 0.04 |
| | 2 (100) | 0.08 | 0.07 | 0.08 | 0.06 | 0.17 | 0.16 | 0.17 | 0.04 |
| *HXT2* | 1 (100) | 0.070 | 0.068 | 0.072 | 0.04 | 0.14 | 0.13 | 0.14 | 0.04 |
| | 2 (102) | 0.096 | 0.08 | 0.11 | 0.05 | 0.13 | 0.13 | 0.14 | 0.06 |

**Supplemental Table 11. Relates to STAR Methods and Key Resources.** Primers for mRNA template amplification from yeast genomic DNA.

| mRNA | Forward Primer | Reverse Primer |
|---|---|---|
| *JJJ1* | TAATACGACTCACTATAGGCAAGAGTTA ATATCAGTATAGTGATATCCTACTACTTC | TTTGTATAGTTGCTTTTCTGGCCTAGTAG AAAGGAAAACTAC |
| *NCE102* | TAATACGACTCACTATAGGATTGATCAA GAAAAAATACAATTGAAAAGG | TAGACAAAGAGAAGGATACTTTAGTTAG ATATAAGAATAGTATGAAAAG |
| *HXT2* | TAATACGACTCACTATAGGACAACAAATT AAATTACAAAAAGAC | TGAGTGGTACAAATAAAAACATCATTAAA AATCG |
| *HSP30* | TAATACGACTCACTATAGGTTATTTATAA TTACAAAAACAAAACAACAAGTTTG | ATAATTTATTTTGCATATTTCATAAAGAAA TTAAAATTAGATTATTAAATATTAAGTTTC |
| *MIM1* | TAATACGACTCACTATAGGACGAAACTG CCACAAGACAGAAATATG | TACGTATAATCTACCGTATGTGTGTGTG TATTTATTTATGTAGGTTGCTAATGC |
| *POP5* | TAATACGACTCACTATAGGCGATAAAAT GGTACGTTTAAAAAGTAGATATATC | TTGTGCTTGAGTTGTATTATAGATAAAAA AATAAAACTACTGCATATACGTGTACATC |
| *SSA1* | TAATACGACTCACTATAGGGCACATCAA AAGAAAAGTAATC | TCTATAGCGTATAGAATATATGTTACATG TATATATATATATAAAGTAAAAACGTTCG G |

### Supplemental Discussion

#### D1. Accuracy of association rate constants from smFRET experiment.

Our mRNA 5'/3' dual-labeling experiments (Fig. S2) indicate that the mRNA ends are within FRET distance for at least 70% of the observation time on mRNAs of variable length. In principle, eIF4E-binding events that occur during the remaining <30% of the observation time might be "missed" under these conditions. However, a dual illumination experiment that allowed simulatenous observation of eIF4E–mRNA binding events that do and do not show FRET, measured previously with a population of ~100 native yeast mRNA molecules (Çetin *et al.*, 2020) revealed that most eIF4E binding events (conservatively, ≥ 80%) that are not attributable to nonspecific surface interaction are actually detected by FRET.

Occasional missed events due to the mRNA ends briefly moving out of FRET range will add one entry for an artificially long apparent arrival time to the arrival-time distribution. However, because we observe many events while the mRNA ends are within FRET range, these small changes to the distribution insignificantly alter fitting of the observed rate constants.
